# Microbiota-dependent indole production is required for the development of collagen-induced arthritis

**DOI:** 10.1101/2023.10.13.561693

**Authors:** Brenda J. Seymour, Brandon Trent, Brendan Allen, Adam J. Berlinberg, Jimmy Tangchittsumran, Widian K. Jubair, Meagan E. Chriswell, Sucai Liu, Alfredo Ornelas, Andrew Stahly, Erica E. Alexeev, Alexander S. Dowdell, Sunny L. Sneed, Sabrina Fechtner, Jennifer M. Kofonow, Charles E. Robertson, Stephanie M. Dillon, Cara C. Wilson, Robert M. Anthony, Daniel N. Frank, Sean P. Colgan, Kristine A. Kuhn

## Abstract

Altered tryptophan catabolism has been identified in inflammatory diseases like rheumatoid arthritis (RA) and spondyloarthritis (SpA), but the causal mechanisms linking tryptophan metabolites to disease are unknown. Using the collagen-induced arthritis (CIA) model we identify alterations in tryptophan metabolism, and specifically indole, that correlate with disease. We demonstrate that both bacteria and dietary tryptophan are required for disease, and indole supplementation is sufficient to induce disease in their absence. When mice with CIA on a low-tryptophan diet were supplemented with indole, we observed significant increases in serum IL-6, TNF, and IL-1β; splenic RORγt+CD4+ T cells and ex vivo collagen-stimulated IL-17 production; and a pattern of anti-collagen antibody isotype switching and glycosylation that corresponded with increased complement fixation. IL-23 neutralization reduced disease severity in indole-induced CIA. Finally, exposure of human colon lymphocytes to indole increased expression of genes involved in IL-17 signaling and plasma cell activation. Altogether, we propose a mechanism by which intestinal dysbiosis during inflammatory arthritis results in altered tryptophan catabolism, leading to indole stimulation of arthritis development. Blockade of indole generation may present a novel therapeutic pathway for RA and SpA.

## Introduction

Rheumatoid arthritis (RA) and spondyloarthritis (SpA) are two forms of inflammatory arthritis characterized by progressive joint inflammation leading to articular destruction. While there are genetic associations of MHC-associated and non-MHC risk alleles in both RA and SpA, these alone are not predictive of disease [1, 2]. Environmental factors such as smoking have been associated with increased risk for RA, and may provide an important trigger in genetically susceptible individuals [3]. Identification of these specific environmental triggers, as well as the mechanisms by which they induce inflammation, may allow for prevention or early treatment of disease by removal of the trigger or specific blockade of the affected pathway.

Evidence of intestinal microbial dysbiosis in both RA [4-8] and SpA [9, 10], as well as significant clinical overlap between SpA and bowel inflammation, suggests that specific microbes or microbial products may serve as an environmental trigger. In RA, this has led to the mucosal origins hypothesis that immune dysregulation in RA begins at mucosal sites [11-13]. RA has a pre-clinical period of 5-10 years preceding clinically apparent disease characterized by intestinal dysbiosis, elevated inflammatory markers, circulating IgA plasmablasts, and autoantibody production at mucosal sites [11, 14], all of which point towards mucosal tissues as the origin of immune dysregulation. Recent identification of an arthritogenic strain of *Subdoligranulum* in individuals at-risk for RA provides further evidence of a mucosal trigger [15]. In SpA nearly 50% of patients have intestinal inflammation at the histologic level [16], and numerous studies support a gut-joint axis in SpA, in which immune dysregulation is initiated in the gut and then spreads systemically to the joint [17]. Despite strong associative evidence of a mucosal origins hypothesis of RA and the gut-joint axis in SpA, the specific mechanisms by which microbes or microbial products trigger intestinal immune dysregulation in the context of inflammatory arthritis have not been elucidated.

We previously demonstrated a progressive gut dysbiosis in the murine model of collagen-induced arthritis (CIA) [32], in which DBA/1 mice are immunized with type-II collagen (CII) emulsified in Complete Freund’s Adjuvant (CFA) at days 0 and 21. In this model, dysbiosis is first observed between days 7-14, prior to the onset of clinically apparent disease, and progresses through day 35 [32], paralleling a microbiome-dependent increase in Th17 cytokines such as IL-17A, IL-22, and IL-23 [32, 33]. Treatment with broad spectrum antibiotics from days 21-35 resulted in significantly reduced CIA severity, suggesting a requirement for the microbiome in the development of disease. Surprisingly, antibiotic treatment did not significantly reduce collagen-specific autoantibody production, but rather impaired complement activation by collagen-specific autoantibodies, which may have been due to altered antibody glycosylation. Together, these findings suggest a key role for the microbiome in shaping the immune response in CIA.

However, the mechanism(s) by which the microbiome drives these inflammatory processes have not been identified. Observations of altered bacterial metabolomes have been described in both murine models of autoimmunity and in patients with autoimmunity [35-39], leading us to hypothesize that certain (yet unidentified) intestinal microbial metabolite(s) would play a key role in the development of CIA. We identified alterations in Tryptophan (Trp) metabolism in CIA in a microbiome-dependent manner, with a significant increase in indole (a bacterial-derived Trp metabolite) production in mice with CIA compared to unimmunized mice and antibiotic-protected CIA mice. We then tested whether indole supplementation was sufficient to induce CIA. We showed through three different methods that indole is indeed required for the development of CIA, and we characterized the cytokine, antibody, and cellular responses affected by indole treatment. Stimulation of human colon lamina propria mononuclear cells (LPMCs) with indole revealed upregulation of pathways similar to those observed in the mice, providing proof of concept that the observed effects of indole in CIA may be relevant to human disease. Altogether, our findings indicate that microbial-derived indole is required for the development of collagen-induced arthritis through enhanced Th17 immunity and modulation of CII-reactive autoantibodies.

## Results

### Microbiome-dependent changes in Trp metabolism are associated with the development of CIA

We previously reported bacterial dysbiosis and mucosal inflammation occurring prior to the onset of arthritis in the CIA model, and that depletion of the microbiome through use of broad-spectrum antibiotics (ampicillin, vancomycin, metronidazole, and neomycin) resulted in a ∼90% reduction in disease severity [32]. To further query the effect of microbial dysbiosis on CIA, and because microbial metabolites can have profound effects on host immunity [40-42], we hypothesized that dysbiosis during CIA would alter the gut metabolome. Thus, we assessed a broad array of central energy and redox metabolites, yielding 244 named metabolites, by liquid chromatography with tandem mass spectrometry (LC-MS/MS) from the cecal contents of DBA/1J mice with CIA and compared them to untreated DBA/1J mice and mice treated with broad-spectrum antibiotics starting at CIA day 21 (CIA+Abx). Partial least squares-discriminant analysis (PLSDA) revealed a distinct, microbiome-dependent metabolome in mice with CIA (Figure 1A). Volcano plot analysis identified several significant differences in Trp metabolites (5-hydroxyindoleacetate, picolinic acid, indoxyl, indolepyruvate, and L-Trp) in the CIA group compared to CIA+Abx, suggesting that CIA induces microbiome-dependent changes in Trp metabolism (Figure 1B).

**Figure 1:**
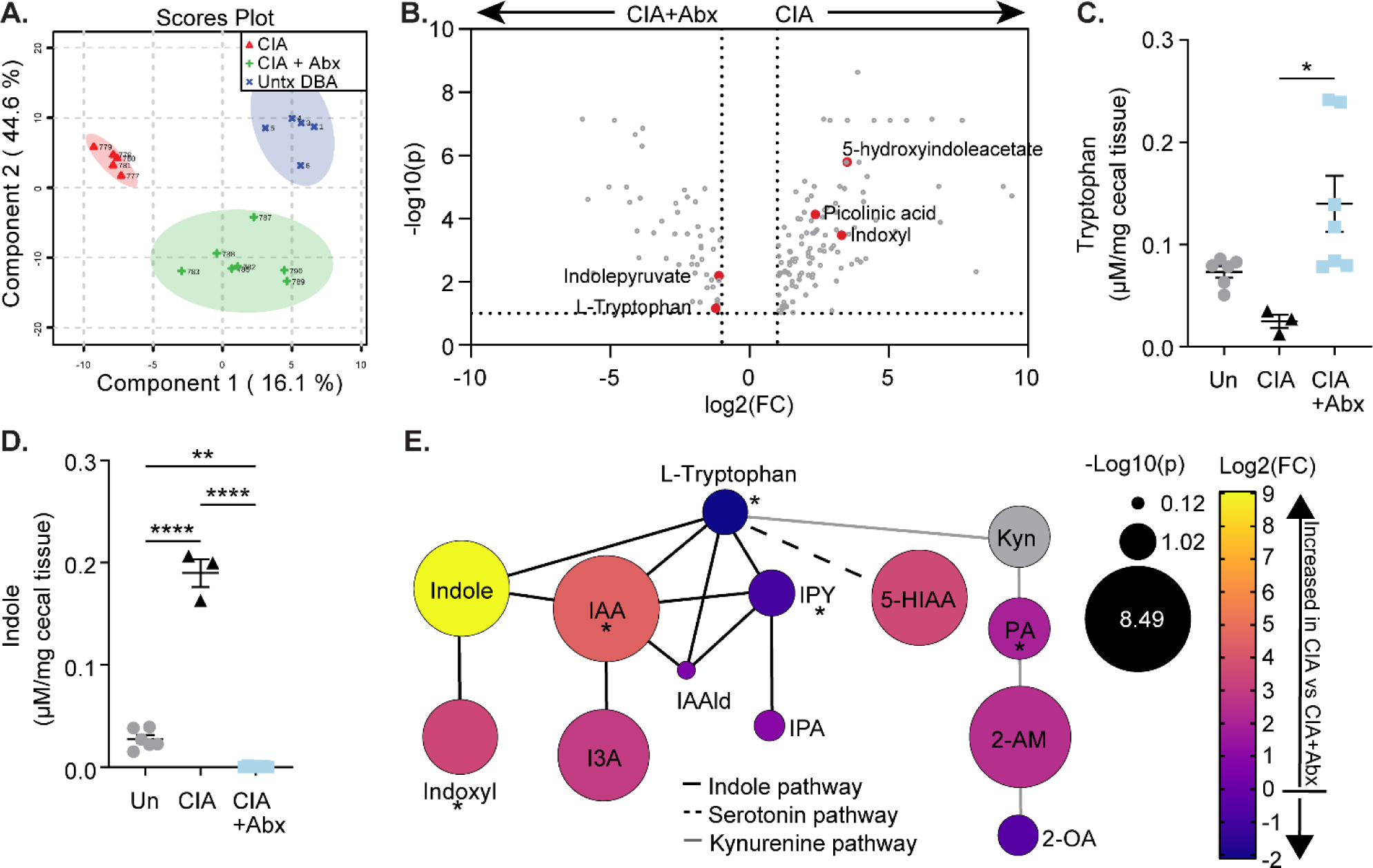
Intestinal metabolomics profiling identifies microbiome-mediated alterations in the tryptophan pathway in mice with CIA. CIA was induced in male 6-week-old DBA/1J mice and cecal contents were harvested at day 35 from mice with CIA (n=3-5), mice with CIA and depleted from microbiota by antibiotic administration after day 21 (CIA+Abx, n=7), or untreated DBA/1J mice (Un, n=5). **(A-B)** LC-MS/MS was used to screen 244 metabolites. **(A)** PLSDA plot of CIA vs CIA+Abx vs Un. **(B)** Volcano plot of CIA+Abx (left) vs CIA (right). **(C-D)** HPLC was used to quantify Trp pathway metabolites indicated on the y-axis in µM. All data were reported as individual mice (symbols) and mean ±SEM (bars) after normalization to weight (mg) of cecal contents. *, p<0.05; ***, p<0.001; ****, p<0.0001 as determined one-way ANOVA with Tukey’s multiple comparisons test. **(E)** Graphical representation of Trp metabolism pathways, showing Trp metabolites identified in the LC-MS/MS analysis (A-B) and HPLC analysis (C-D). Log2(fold change) was calculated for CIA vs CIA+Abx and is represented in color gradient from yellow (more increased in CIA) to blue (more increased in CIA+Abx). The size of each circle represents -log10(pvalue) of an unpaired t-test between CIA vs CIA+Abx. IAA, indole-3-acetic acid; I3A, indole-3-carboxaldehyde; IPY, indolepyruvate; IPA, indolepropionic acid; 5-HIAA, 5-hydroxyindoleacetate; Kyn, L-kynurenine; IAAld, indole-3-acetaldehyde; PA, picolinic acid; 2-AM, 2-aminomuconate; 2-OA, 2-oxoadipate. Asterisk (*) denotes trends in metabolites that were also observed in *Isolate 7*-colonized mice. Line color denotes pathway: black, indole; dashed, serotonin; grey, kynurenine.

Targeted HPLC quantification of seven Trp metabolites from cecal contents revealed significant decreases in cecal Trp in the CIA group compared to CIA+Abx, along with a significant increase in indole (Figure 1C and D), suggesting that microbiome-mediated Trp metabolism is skewed towards indole production in CIA. Decreased Trp in mice with CIA compared to CIA+Abx is consistent with microbial consumption of Trp, as reported previously [35]. Trp is generally metabolized by three pathways: indole (primarily microbial), kynurenine, and serotonin (both of which are primarily host pathways). To better asses which Trp pathways were affected in CIA, Trp metabolites that were identified in either the LC-MS/MS screen (Figure 1A-B) or HPLC (Figure 1C-D, and Supplemental Figure 1A-D) were then manually mapped (Figure 1E) using the TrpNet.ca database [54] for reference, which provides pathway/relationship information on 108 Trp metabolites identified in the mouse and human microbiome. Of the metabolites screened in the indole pathway, 4/7 (indole-3-acetic acid (IAA), indole, indole-3-carboxaldehyde (I3A), and indoxyl) were significantly increased in CIA compared to CIA+Abx (p<5.9E5), 2/7 were increased but not statistically significant (indolepropionic acid (IPA) and indole-3-acetaldehyde (IAAld)), and 1/7 was decreased (indolepyruvate (IPY)). Of these metabolites, indole, I3A, IPA, and IPY are produced exclusively by the microbiome, while IAA, indoxyl, and IAAld can be produced by both host and microbiome[54]. While metabolites from the serotonin and kynurenine pathways were also significantly increased (5-hydroxyindoleacetate (5-HIAA), picolinic acid (PA), and 2-aminomuconate (2-AM)), the largest differences were observed within the indole pathway. Furthermore, 5HIAA and PA are produced exclusively by host, not microbiome [54]. Because of the requirement for the microbiome in the development of CIA, we focused our attention on microbial-derived metabolites within the indole pathway, and in particular, indole, which had the highest differential abundance: a 528-fold increase in mice with CIA compared to the CIA+Abx group, and an 8-fold increase in mice with CIA compared to untreated DBAs (Figure 1D-E).

Because of the observed alterations in Trp metabolism in CIA, as well as our previous observations of altered Trp in SpA [9] and recent reports in murine lupus and experimental autoimmune encephalitis (EAE) models [35, 48], we queried whether Trp metabolism was altered in another murine model of inflammatory arthritis. We recently identified an arthritogenic bacterium *S. didolesgii Isolate 7*, isolated from the stool of a patient at-risk for RA. Mono-colonization of germ-free mice with *Isolate 7* stimulates spontaneous joint swelling, Th17 cell expansion, and production of autoantibodies in a pattern that mimics early RA [15]. Using unbiased LC-MS/MS screening of metabolites from cecal contents of *Isolate 7-*colonized mice, compared to non-arthritogenic *S. didolesgii Isolate 1-*colonized mice, we observed a pattern of altered Trp metabolism similar to mice with CIA, with significant increases in IAA, indoxyl, and PA, and decreases in IPY and L-tryptophan (Supplemental Figure 1E and asterisks in Figure 1E).

We then sought to identify the microbial source(s) of indole in CIA using paired 16S amplicon sequencing of fecal contents with cecal metabolomics by LC-MS/MS. Bacterial taxa belonging to the phyla Firmicutes (*Lactobacillales*, *Hydrogenoanaerobacterim*) and Bacteroidetes (*Barnesiella*, *Rikenellaceae*, *VC2.1-Bac22*) showed strong correlation with indoxyl levels by Spearman correlation (Supplemental Figure 1F). Paired indole-16S data was not available, so indoxyl was used as a proxy in the Spearman correlation (indole is metabolized by hepatic CYP450 enzymes into indoxyl [55]). Of the taxa that correlated with indoxyl, all were elevated in CIA compared to CIA+Abx (Supplemental Figure 1G), indicating that their expansion may result in greater indole production. Finally, we found that indole levels significantly correlated with disease severity in mice with CIA (Supplemental Figure 1H). These results, in addition to observations of altered Trp metabolism in patients with SpA and RA [9, 47, 56-59], prompted us to further interrogate the role of bacterial-derived Trp metabolites in inflammatory arthritis.

### Bacterial Trp catabolism to generate indole is required for the development of inflammatory arthritis in CIA

Next, we hypothesized that dysbiosis following CIA induction leads to increased indole production, which then enhances disease severity. We first tested if indole could reverse the disease protection afforded by antibiotic-induced ablation of microbiota during CIA [32]. As done previously [32], CIA was induced by immunization of bovine type II collagen in CFA on days 0 and 21. At day 21, antibiotics including ampicillin, metronidazole, neomycin, and vancomycin were added to the drinking water. In addition, 0.1 mg/ml indole was included in the drinking water for one group of mice at this time. We observed a ∼3-fold increase in arthritis severity 35 days following the initial immunization when adding indole in the drinking water of CIA+Abx mice (Figure 2A), suggesting that indole was sufficient to, at least partially, replace the role of the bacterial microbiome during CIA.

**Figure 2.**
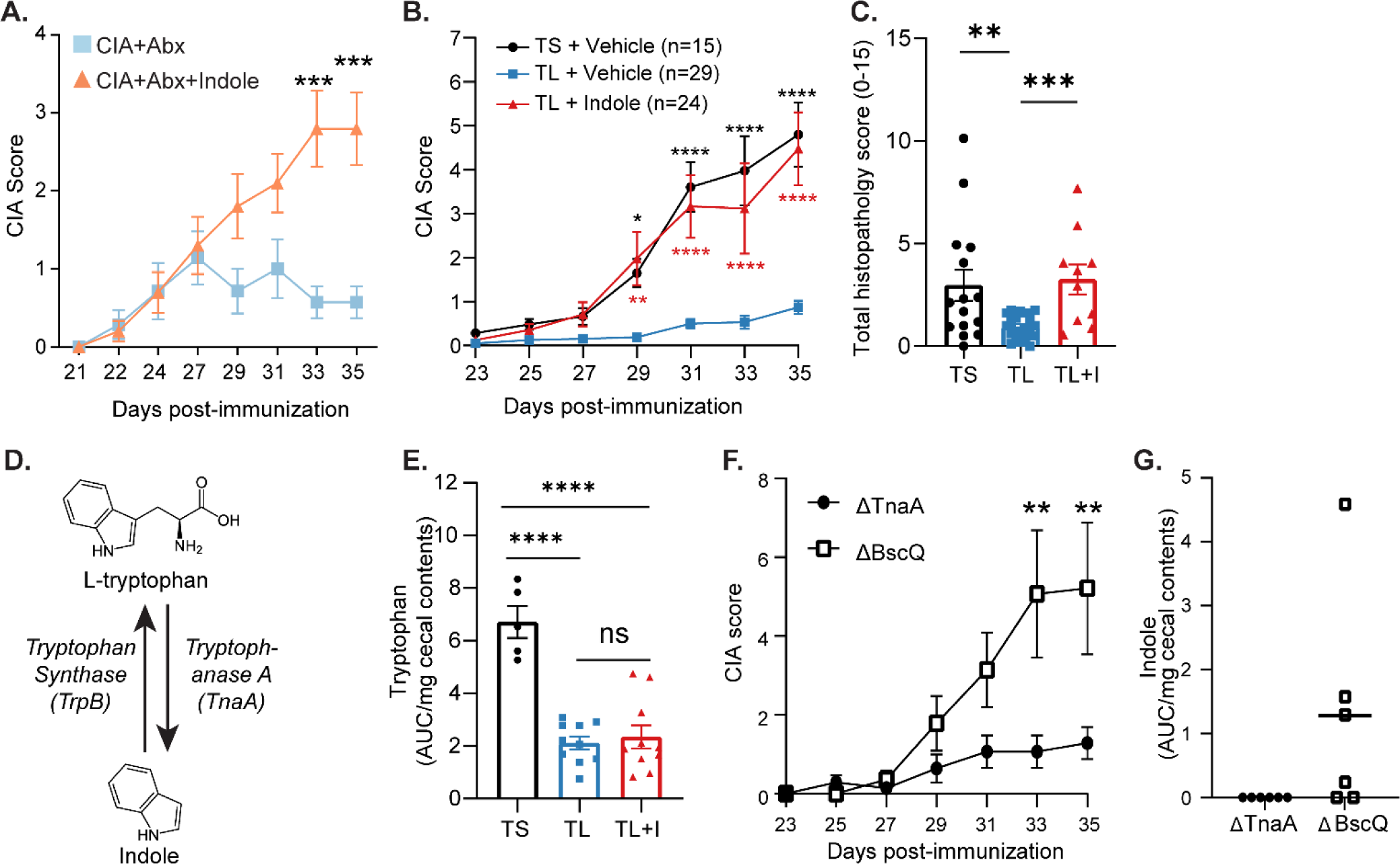
Indole is required for CIA. **(A)** CIA was induced in male 6-week-old DBA/1J mice. On days 21-35, mice were treated with antibiotics ± 0.1 mg/ml indole in the drinking water. N=10 (Abx+Indole); N=7 (CIA+Abx). **(B)** Male 6-week-old DBA/1J mice were fed a TL or TS diet starting at day -1 through the duration of the experiment. Following induction of CIA, mice were treated with indole (200µl of a 10mM solution) or vehicle control (0.33% methanol) by oral gavage every other day starting on day 0. Arthritis scores were assessed every other day from day 21-35. N=29 (TL+Vehicle); N=24 (TL+Indole); N=15 (TS+Vehicle) pooled from three independent experiments. Red asterisks: TL+Indole vs TL+Vehicle; black asterisks: TS+Vehicle vs TL+Vehicle. TS+Vehicle vs TL+Indole was not statistically significant. **(C)** The sum of the inflammation, pannus, and bone erosion score of H&E stained paws is plotted as the total histology score (maximum score of 15). N=10-20 from two independent experiments. **(D)** Schematic of Trp breakdown into indole by bacterial *Tryptophanase A*, and Trp synthesis from indole by bacterial *Tryptophan synthase*. **(E)** HPLC analysis of Trp in cecal contents from mice with CIA at day 35, plotted as area under the curve (AUC), normalized to weight (mg) of cecal contents. N=5-10 from one experiment. **(F)** Male 6-week-old germ-free DBA/1 mice were colonized with *E. coli BW25113* mutants (*ΔtnaA* or *ΔBcsQ*) with 10^8^ CFU by oral gavage on day -7 before CIA induction. N=7 per group. **(G)** HPLC analysis of indole in cecal contents from CIA mice colonized with either *ΔtnaA* or *ΔBcsQ* at CIA day 35. Indole levels were plotted as AUC/mg cecal content weight. Data are reported as mean ± SEM. *, p<0.05; **, p<0.01; ***, p<0.001; ****, p < 0.0001 as determined by two-way ANOVA with Bonferroni adjustment for multiple comparisons (A, B, F) and unpaired t-test (C, E, G).

We then tested the effects of various levels of dietary Trp on CIA. Because Trp is an essential amino acid, mice fed a Trp-deficient (0% Trp) diet lose ∼15-20% body weight within 7 days [35]. Therefore, similar to previous studies [35], the Trp-deficient diet was replaced with a Trp-sufficient (TS) diet, which contains 0.18% L-Trp, on the weekends for a cumulative Trp-low (TL) diet (0.05% L-Trp). With this TL diet, mice did not lose more than 15% of their starting body weight (Supplemental Figure 2A). To ensure consistent dosing, indole (10mM) or vehicle (0.33% methanol) were added back by oral gavage every other day starting on day 0 for the duration of the experiment. The TL+Vehicle group had a significant reduction in disease severity and incidence, with an average CIA score of ∼1 (Figure 2B) and 55% CIA incidence (Supplemental Figure 2B), suggesting a requirement for dietary Trp in the development of CIA. Adding back indole resulted in a 4.5-fold increase in CIA severity compared to TL+Vehicle mice (Figure 2B) with 92% CIA incidence (Supplemental Figure 2B). There was no significant difference in CIA scores or incidence in the TL+Indole group compared to TS+Vehicle, suggesting that indole supplementation fulfills the requirement for dietary Trp in the development of CIA. Evaluation of pathology by H&E staining in a subset of mice revealed a significant decrease in inflammatory infiltrate, pannus, and bone resorption in the TL+Vehicle group compared to TS+Vehicle, which was completely restored with indole supplementation (Figure 2C and Supplemental Figure 2C-G). Across all groups, the histologic inflammation score significantly correlated with the macroscopic arthritis score at day 35 (Supplemental Figure 2H).

Trp is an essential amino acid in mammals and can be synthesized from indole by microbial *Tryptophan synthase* (*TrpB*) [60] (Figure 2D). As such, we investigated whether the reversal of TL-mediated protection in CIA with indole supplementation is due to restoration of Trp levels by *TrpB*. HPLC analysis of cecal metabolites at CIA day 35 did not show a significant difference in Trp levels between TL+Vehicle and TL+Indole mice, and both groups had significantly lower cecal Trp levels than TS+Vehicle mice (Figure 2E), suggesting that indole was not acting as a substrate for Trp synthesis. These data suggest that the differences in disease severity between TL+Vehicle and TL+Indole mice cannot be explained by Trp availability, and rather, likely is due to an effect of indole on either the host or the microbiome.

To further query the specific requirement for indole in CIA, we targeted microbial *Tryptophanase A* (*TnaA),* which is the sole enzyme responsible for indole production [61, 62] (Figure 2D). Germ-free DBA/1 mice were colonized with 10^8^ CFU of either an *E. coli* K12 strain (BW25113 *ΔtnaA)*, which is unable to produce indole due to deletion of *Tryptophanase A* [63], or an isogenic control (*E. coli* BW25113 *ΔbcsQ*), which produces normal levels of indole. Mice were maintained on standard rodent chow containing 0.2% L-Trp. Stable colonization of both *ΔtnaA* and *ΔbcsQ* was confirmed by qPCR using global rpoB primers (Supplemental Figure 2I). Mice colonized with *ΔtnaA* had a significant ∼5-fold reduction in CIA severity compared to the *ΔbcsQ* control group (Figure 2F). HPLC analysis of cecal contents at day 35 confirmed that indole was only produced in the *ΔbcsQ* controls (Figure 2G). Together, these suggest that indole produced by microbial *Tryptophanase A* is required for the development of CIA.

### Indole minimally impacts bacterial dysbiosis due to a low Trp diet

We next sought to identify the mechanism through which indole incites disease. While we hypothesized that indole has a direct immunogenic effect on the host, leading to the development of CIA, an alternate hypothesis is that indole indirectly affected disease outcomes through its effects on the microbiome [64], as previous studies have shown that fecal transfer of “arthritogenic” or “non-arthritogenic” microbiomes predicts CIA severity and susceptibility in recipient mice [33].16S rRNA gene sequencing of fecal contents revealed alpha diversity was lowest in the TS+Vehicle group (Figure 3, A-C). Interestingly, while the TL+Indole group developed similar levels of arthritis severity as the TS+Vehicle group, the TL+Indole group maintained higher alpha diversity (Figure 3, A-C). Beta diversity was most significantly affected by the TL diet as the TS+Vehicle group showed greater dissimilarity from the TL+Vehicle and TL+Indole groups by PCA (Figure 3D and Supplemental Figure 3A) and PERMANOVA (Figure 3E and Supplemental Figure 3B). We then identified individual bacterial taxa driving the observed differences in microbial diversity. Ten taxa were expanded in the TS+Vehicle group compared with TL+Vehicle (FDR-corrected p<0.05; Figure 3F-G), including several that we have previously shown to be expanded in CIA (e.g., *Lachnospiraceae*) [32]. Conversely, *Turicibacter, Clostridium, and Peptostreptococcaceae* were significantly elevated in the TL+Vehicle group compared to TS+Vehicle (FDR-corrected p<0.05; Figure 3, F-G). These three genera also accounted for the largest effect sizes in the comparison of TS+Vehicle vs. TL+Indole groups (FDR-corrected p<0.05; Figure 3H-I). No taxa were found to be differentially abundant between TL+Vehicle and TL+Indole groups (n=10 per group) after adjusting p-values for multiple comparisons (i.e., all FDR-corrected p values were >0.05) (Supplemental Figure 3C-D). Taken together, the results of microbiome profiling suggest that dietary tryptophan significantly impacts fecal bacterial communities, but indole supplementation only minimally altered the diversity and composition of the fecal microbiota, and likely does not explain the arthritis phenotype observed following indole supplementation.

**Figure 3.**
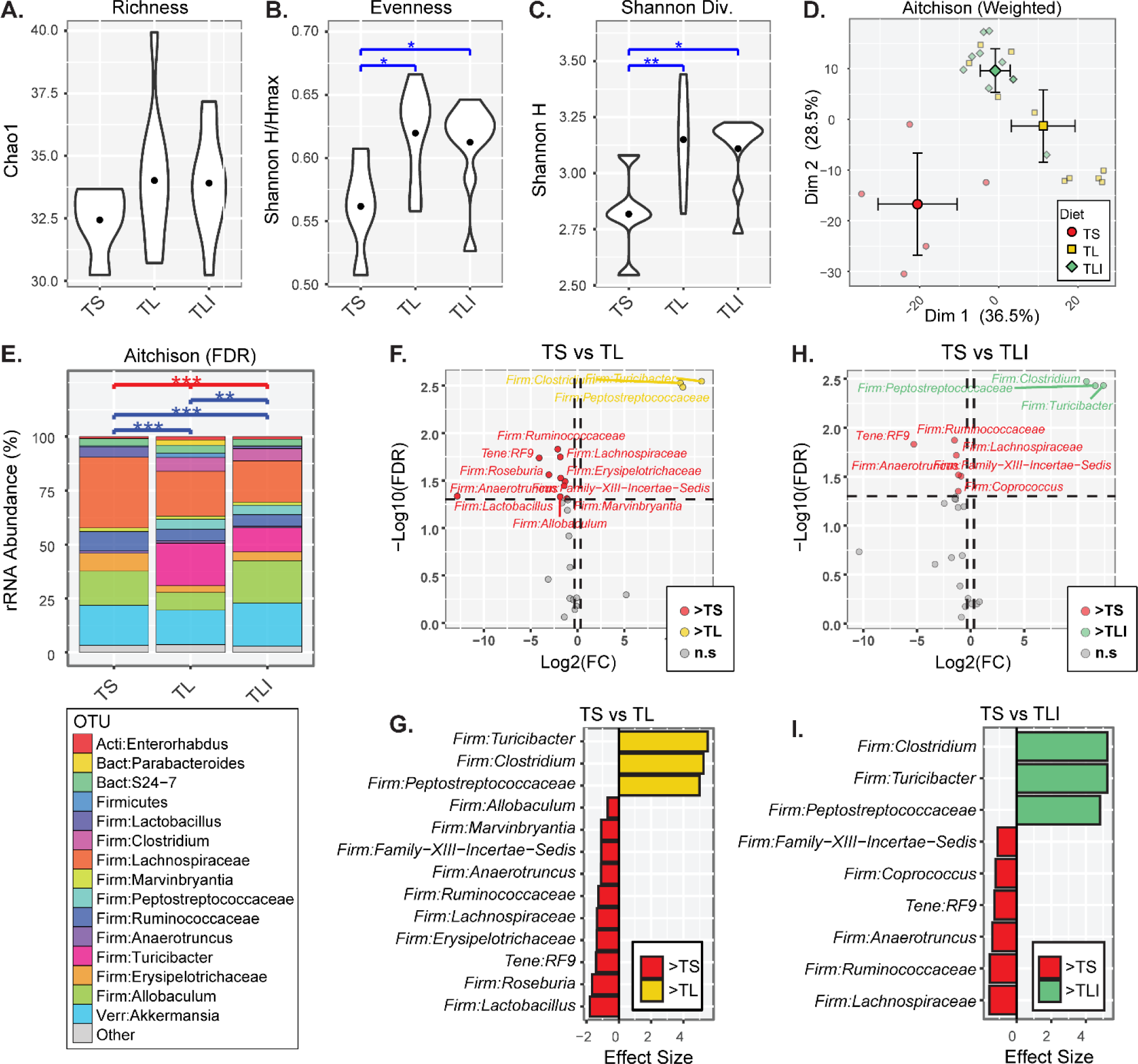
Indole minimally affects bacterial dysbiosis imparted by a TL diet during CIA. Male 6-week-old DBA/1J mice were fed a TL or TS diet starting at day -1 through the duration of the experiment. CIA was induced and indole (200ul of a 10mM solution) or vehicle control (0.33% methanol) was added by oral gavage every other day starting on day 0. On day 35, fecal pellets were harvested, genomic DNA isolated, and 16S rRNA sequencing was utilized to assess microbial diversity. **(A-C)** Alpha diversity indices are shown for each group (TS = TS+Vehicle (n=5); TL = TL+Vehicle (n=10); TLI = TL+Indole (n=10)). Differences between groups were assessed by ANOVA: *, p<0.05 and **,p<0.01. **(D)** PCoA in which smaller, lighter symbols represent individual mice and large, darker symbols represent group means + 95% confidence intervals for PC1 and PC2. **(E)** Bar charts showing mean distributions of taxa for each group. Taxa with relative abundances <1.0% were collapsed into the “Other” category to simplify the figure. Differences in beta-diversity between groups were assessed using PERMANOVA tests with the weighted Aitchison dissimilarity index: **, p<0.01; ***, p<0.001; and ****, p<0.0001. **(F-I)** Volcano and effect size plots generated by ANOVA-like differential expression (ALDEx2) analysis indicate taxa that were significantly enriched or depleted (FDR-corrected p-value < 0.05) in mice with CIA on one diet compared to another: TS+Vehicle vs TL+Vehicle **(F-G)**, TL+Vehicle vs TL+Indole **(H-I).**

### Indole induces a unique inflammatory cytokine signature in CIA

Because the effects of indole on the fecal microbiota were minimal in the setting of a TL diet, we then assessed the effect of indole on immune cell function in CIA. First, because pro-inflammatory cytokines are key mediators of disease in CIA, RA, and SpA, we assessed the effects of the TL diet and indole supplementation on serum cytokine production throughout the course of CIA. Terminal serum was collected from mice with CIA at days 14, 21, and at the plateau of disease severity (day 35-50, depending on the experiment). To account for experiment-to-experiment variability, serum cytokine concentrations were normalized to the mean of the TS+Vehicle group for each experiment/timepoint to allow for better group-to-group comparisons at each timepoint. Cytokine concentrations of the TS+Vehicle group (which were used for normalization) are shown in Supplemental Figure 4 (A-F) compared to naïve (unimmunized) DBA/1 mice.

Except for a small, but significant, difference in IL-1β at CIA day 14 in the TL+Indole group compared to TL+Vehicle (Figure 4A), there were no significant differences between groups in any of the cytokines measured at CIA days 14 and 21 (prior to arthritis onset). However, by the plateau of disease (CIA day 35-50), the TL+Vehicle diet resulted in a significant reduction in IL-1β and IL-6, with significantly increased IL-10 compared to mice on the TS+Vehicle diet at CIA day 35+, but not at earlier timepoints (days 14 and 21) (Figure 4A-C). These findings are consistent with the requirement for both IL-6 and IL-1β in the development of CIA, and the protective role of IL-10 [65-67].

**Figure 4.**
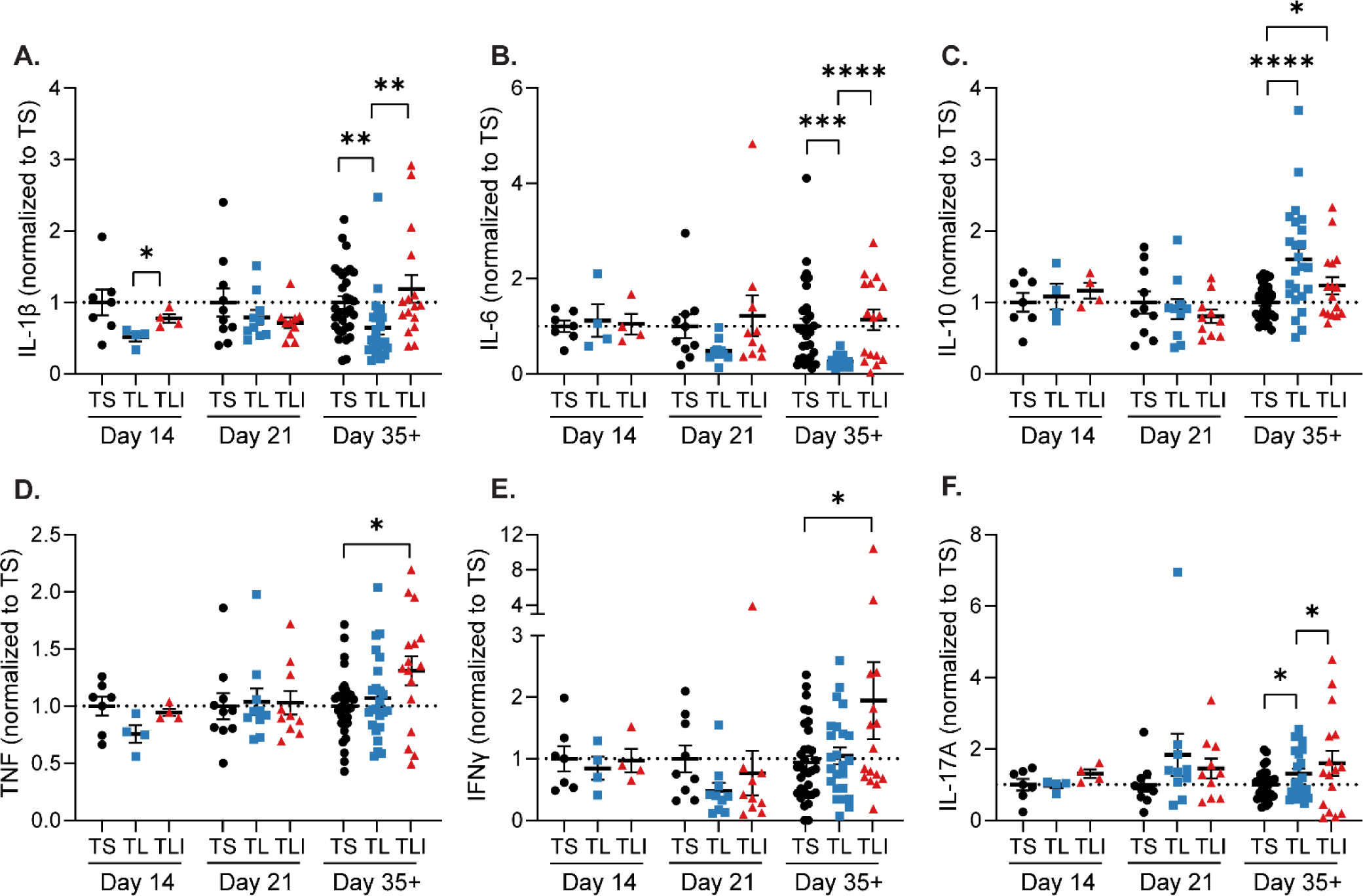
Indole alters the cytokine profile in CIA. Male 6-week-old DBA/1J mice were fed a TL or TS diet starting at day -1 through the duration of the experiment. Following induction of CIA, mice were treated with indole (200µl of a 10mM solution) or vehicle control (0.33% methanol) by oral gavage every other day starting on day 0. **(A-F)** Terminal serum was collected at days 14, 21, and at the plateau of disease (day 35-50) from mice with CIA fed a TL or TS diet and treated with indole (200ul of a 10mM solution) or vehicle control (0.33% methanol) and analyzed by an 8-plex immunoassay (Mesoscale). To account for experiment-to-experiment variability, serum cytokine concentrations (as denoted on the y-axis) were normalized to the mean of the TS+Vehicle group for each experiment/timepoint. N=4-8 (day 14), 10 (day 21), and 16-28 (day 35-50) pooled from nine independent experiments and plotted as individual mice (symbols) and mean ±SEM (bars). *, p<0.05; **, p<0.01; ***, p<0.001 and ****, p<0.0001 by unpaired t-test.

Indole supplementation restored IL-1β and IL-6 levels at day 35+ (Figure 4AB). Surprisingly, despite nearly identical CIA severity in the TL+Indole and TS+Vehicle groups (Figure 2B), indole supplementation did not fully replicate the cytokine pattern observed in TS+Vehicle mice. Rather, indole supplementation created a unique cytokine signature, with significant increases in IL-10, TNF, and IFNγ compared to the TS+Vehicle group (Figure 4C-E). There was a small, but significant, increase in IL-17A in both TL+Vehicle and TL+Indole compared to the TS+Vehicle group (Figure 4F). IL-22, IL-23, and GM-CSF were also measured at day 35 but were either at or below the limit of detection (Supplemental Figure 4G-I). Altogether, these data suggest that indole induces a unique pro-inflammatory signature in CIA.

### Indole alters autoantibody pathogenicity

Another hallmark of CIA is the development of collagen (CII)-specific autoantibodies [68, 69]. As such, we queried whether indole affects production of type-II collagen (CII) antibodies. There were no significant differences in total IgG, anti-CII IgG, or germinal center B cell numbers between groups (Supplemental Figure 5A-C), and anti-CII IgG levels did not correlate with CIA severity (Supplemental Figure 5D), suggesting that the arthritis severity observed in the TL+Indole compared to TL+Vehicle groups cannot be attributed solely to anti-CII autoantibody quantity, in agreement with previous studies [32, 70].

We then hypothesized that indole may affect the function, rather than quantity, of anti-CII antibodies. Indeed, we have previously demonstrated reduced complement fixation by anti-CII antibodies in CIA+Abx mice [32], suggesting that microbial stimuli may alter the ability of anti-CII antibodies to activate complement, thus altering their pathogenicity. In serum from TL+Indole mice compared to TL+Vehicle, we observed a significant increase in complement fixation by anti-CII antibodies (Figure 5A). Complement deposition in the joints followed the same pattern, with a significant decrease in complement deposition in the TL+Vehicle group compared to TS+Vehicle, which was restored with indole supplementation (Figure 5B and Supplemental Figure 5E-F). Furthermore, complement fixation by CII-specific antibodies and C3 deposition in the joint both correlated with CIA severity (Supplemental Figure 5G-H), providing further evidence that antibody function rather than concentration is more important in CIA pathogenesis.

**Figure 5.**
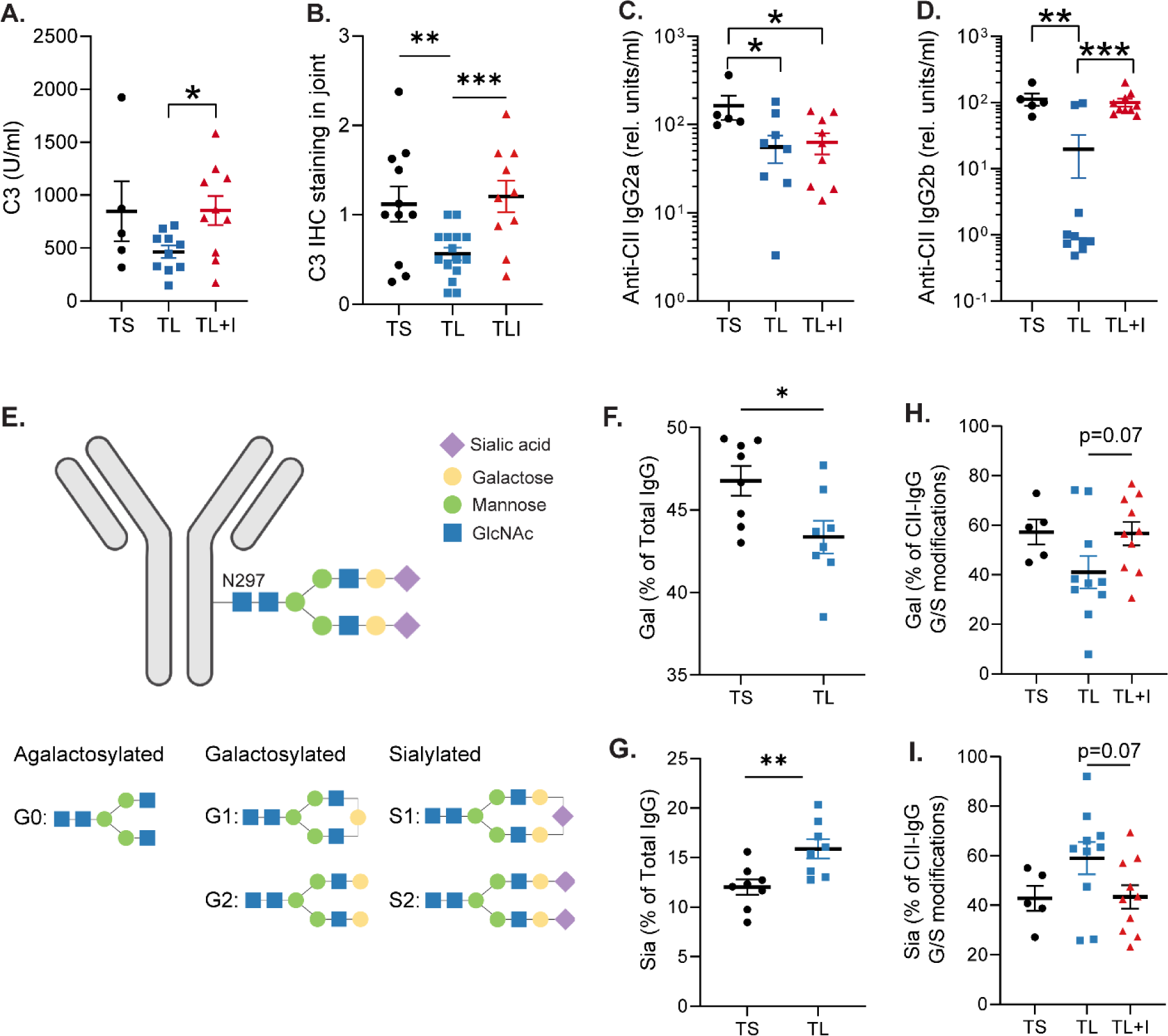
Indole alters complement activation, IgG subclass, and glycosylation. Male 6-week-old DBA/1J mice were fed a TL or TS diet starting at day -1 through the duration of the experiment. Following induction of CIA, mice were treated with indole (200µl of a 10mM solution) or vehicle control (0.33% methanol) by oral gavage every other day starting on day 0. **(A)** Day 35 serum was evaluated by ELISA for C3 binding to anti-CII IgG. N=5-10 from one independent experiment. **(B)** FFPE joints were stained for complement C3 by immunohistochemistry and staining intensity was scored. Each datapoint represents the average complement deposition score of all 4 paws for one mouse (maximum score = 3). N=10-20 per group pooled from two independent experiments. **(C-D)** Day 35 serum was evaluated by ELISA for anti-CII IgG2a **(C)** and anti-CII IgG2b **(D).** N=5-10 per group from one independent experiment. **(E)** Diagram of possible glycosylation patterns on N297 of the IgG Fc domain. Blue circles = N-acetylglucosamine; green circles = mannose; yellow circles = galactose; purple diamonds = sialic acid. **(F-G)** Total IgG was purified from serum and IgG glycosylation patterns were assessed by liquid chromatography with mass spectrometry (LC-MS/MS). % Galactosylation (Gal) and % Sialylation (Sia), were plotted, respectively. Galactosylation and Sialylation were calculated as a % of all glycoforms (G0, G1, G2, S1, and S2). N=10 per group from one independent experiment. **(H-I)** In a separate experiment, anti-CII IgG was purified using CII-linked CNBr Sepharose 4B beads. IgG glycosylation patterns were assessed by LC-MS/MS. Galactosylation and sialylation are potted as % of G1, G2, S1, and S2 glycoforms only. N=5-10 per group from one representative experiment. For all panels, values are plotted as individual mice (symbols) and mean ± SEM (bars). *, p<0.05; **, p<0.01; ***, p<0.001; ****, p < 0.0001 as determined by unpaired t-test.

We then examined anti-CII IgG isotypes and Fc glycosylation, as both have been shown to affect complement binding [71-76]. Whereas we did not observe significant differences in anti-CII IgG1 (Supplemental Figure 6A), the TL+Vehicle treatment significantly reduced anti-CII IgG2a and IgG2b concentrations (Figure 5C-D). Intriguingly, while indole supplementation was not sufficient to rescue anti-CII IgG2a levels, anti-CII IgG2b was restored to the level of the TS+Vehicle group (Figure 5C-D). Furthermore, anti-CII IgG2b levels significantly correlated with CIA severity and complement activation (Supplemental Figure 6B-C), whereas anti-CII IgG1 only weakly correlated and IgG2a had no correlation (Supplemental Figure 6D-E). Because complement-fixing IgG2b has been demonstrated to be highly pathogenic in other models [77], the observed changes in complement activation with indole supplementation may be explained by increased anti-CII IgG2b isotype switching.

We next assessed IgG glycosylation. The N297 residue of the IgG Fc chain is modified by glycans in response to physiological changes. The main IgG-Fc glycoforms are agalactosylated (G0), galactosylated (G1 or G2) or sialylated (S1 or S2) [75, 78] (Figure 5E). IgG-Fc glycosylation patterns are altered in patients with RA, with decreased sialylation compared to controls [79, 80]. N-terminal sialic acid residues have been reported to have anti-inflammatory properties, specifically through the induction of inhibitory FCγRIIb receptors [81], and can act as a “cap” that blocks complement C1q from binding to galactose residues [70, 71, 76]. We first assessed glycosylation of total purified serum IgG from mice with CIA at day 35 by measuring the G0, G1, G2, S1 and S2 glycoforms. The TL+Vehicle group had significantly reduced galactosylated (G1 + G2) and correspondingly increased sialylated (S1 + S2) IgG compared to the TS+Vehicle group (Figure 5F-G). In a second experiment (plotted separately due to HPLC batch effects), indole supplementation resulted in an intermediate level of IgG sialylation (Supplemental Figure 6F-G).

We then reasoned that glycosylation may be more significantly affected in the antigen-specific antibody response, and examined glycosylation of purified CII-specific antibodies from day 35 serum. Due to a low yield of CII-IgG after purification, there was a high level of background in the HPLC analysis that prevented us from quantifying the agalactosylated (G0) glycoform. Therefore, we quantified the relative levels of either galactosylated (G1 + G2) or sialylated (S1 + S2) glycoforms. Although there were no statistically significant differences between groups, we observed the same pattern in the CII-specific fraction as observed with total IgG: the TL+Vehicle group had decreased galactosylation and increased sialylation compared to the TS+Vehicle group (Figure 5H-I). Indole supplementation appeared to restore the glycosylation pattern of the TS group but was not statistically significant (p=0.07) (Figure 5H-I). These data suggest that indole may promote pathogenic antibody glycosylation in an antigen-specific manner. Altogether, these findings suggest that the observed increase in complement activation following indole supplementation may be due to isotype switching and/or antibody glycosylation patterns that favor complement binding.

### Indole may enhance Th17 immunity

Previous studies have shown that the IL-23/IL-17 axis (Th17) but not IL-12/IFNy (Th1) axis is required for CIA[82], and that the IL-23/Th17 axis promotes CIA, at least in part, through altered CII-antibody glycosylation [70]. Based on our observations that indole supplementation leads to decreased antibody sialylation (Figure 5I), we hypothesized that indole supplementation may promote CIA through the IL-23/Th17 axis. As such, we first assessed whether Th17 cells were affected by indole supplementation. At CIA day 35, splenic total CD4+ T cell numbers were not significantly different between the TL+Vehicle and the TS+Vehicle groups, suggesting that reduced dietary Trp in mice with CIA on the TL diet does not alter global T cell numbers (Supplemental Figure 7A-B). We then assessed the proportions of naïve, effector, and central memory CD4+ T cells. The TL+Vehicle diet did not affect the distribution of effector and naïve CD4+ T cells, although there was a slight reduction in central memory CD4+ T cells compared to the TS+Vehicle group (Figure 6A-C). Surprisingly, there was a significant reduction in the proportion of naïve CD4+ T cells in the TL+Indole group, and a corresponding increase in effector and central memory populations compared to the TL+Vehicle group (Figure 6A-C). When assessed by cell number, we observed a significant reduction in total number of naïve CD4+ T cells in the TL+Indole group, but no changes in effector or central memory cell numbers (Supplemental Figure 7C-E). In the colon, there were no significant differences in proportion of naïve, effector, and central memory CD4+ T cells (Supplemental Figure 7F-H).

**Figure 6.**
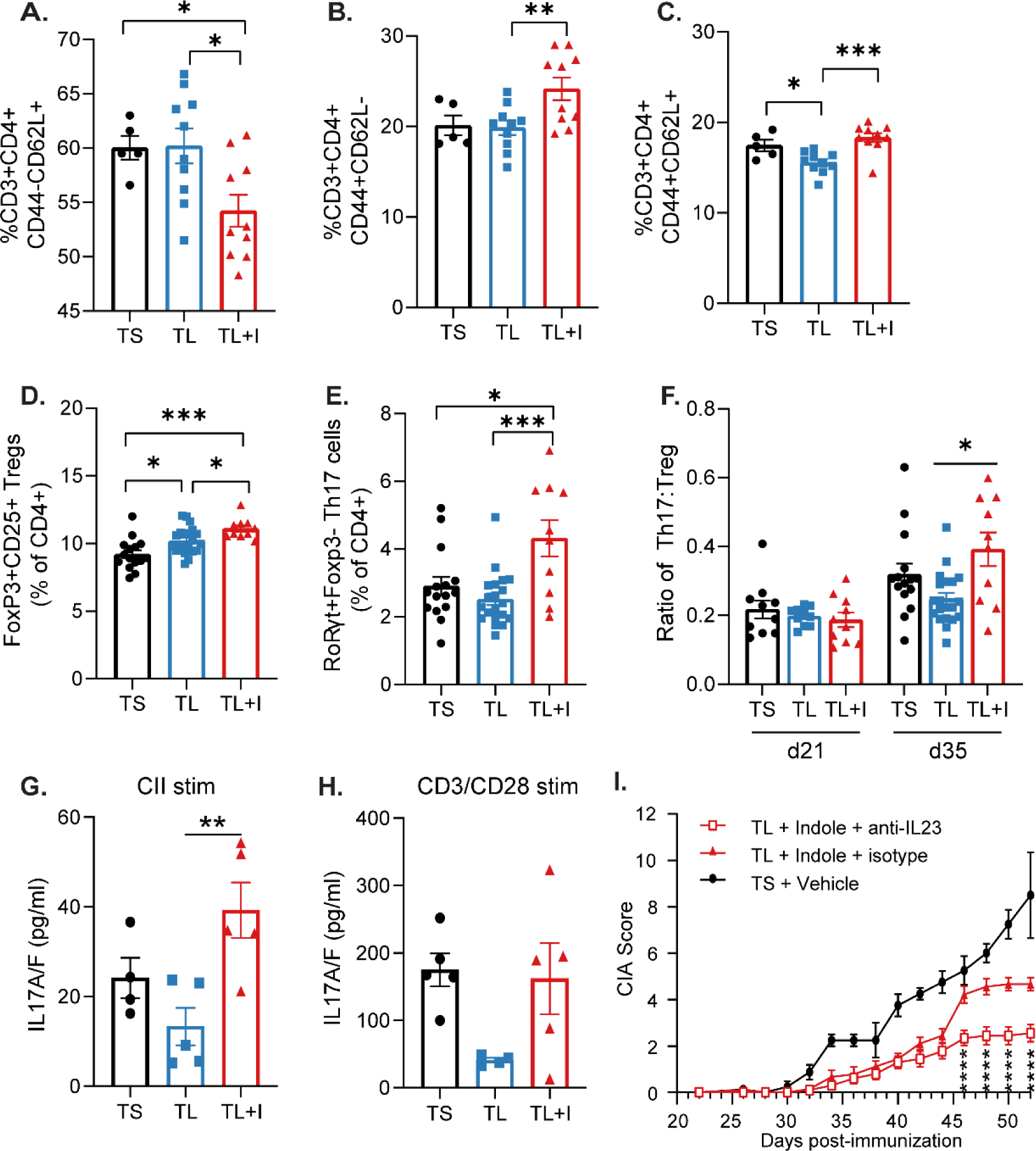
Indole skews towards Th17 cells. Male 6-week-old DBA/1J mice were fed a TL or TS diet starting at day -1 through the duration of the experiment. Following induction of CIA, mice were treated with indole (200µl of a 10mM solution) or vehicle control (0.33% methanol) by oral gavage every other day starting on day 0. Spleens were harvested at day 35 for analysis by flow cytometry. **(A)** Splenic T_naïve_ (CD44-CD62L+) as % of CD4+ T cells. **(B)** Splenic T_effector_ (CD44+CD62L-) as % of CD4+ T cells. **(C)** Splenic T_CM_ (CD44+CD62L+) as % of CD4+ T cells. N=5-10 per group from one independent experiment for panels A-C. **(D)** Splenic FoxP3+RORγt-CD25+ Tregs are plotted as the percent of total CD4+ T cells. **(E)** Splenic CD3+CD4+FoxP3-RORγt+ Th17 cells are plotted as the percent of total CD4+ T cells. **(F)** Ratio of splenic Th17 to Treg cells at CIA days 21 and 35. N=10-20 per group from two independent experiments (day 35) (panels D-F) and n=10 per group (day 21) from one independent experiment (Panel F). **(G-H)** Total splenocytes from CIA day 35 were harvested and re-stimulated with bovine type II collagen **(G)** or CD3/CD28 Dynabeads **(H)**; supernatant was saved and IL17A/F was measured by MSD**. N**=5 per group from one independent experiment. **(I)** TL+Indole treated mice received IP injections of 100µg of anti-IL23p19 or isotype (anti-HRP) on CIA days 0, 7, 14, and 21 and CIA severity monitored. N=10 (TL+Indole+anti-IL-23), N=10 (TL+Indole+Isotype), and N=5 (TS+Vehicle). Asterisks show the comparison between TL+Indole+Isotype vs TL+Indole+anti-IL23. For all panels, values are plotted as individual mice (symbols) and mean ± SEM (bars). *, p<0.05; **, p<0.01; ***, p<0.001; ****, p < 0.0001 as determined by unpaired t-test (panels A-H) or two-way ANOVA with Bonferroni adjustment for multiple comparisons (panel I).

Because of the skewed proportions of naïve and effector CD4+ T cells observed in the spleen, we next assessed effector CD4+ subsets, including Foxp3+ Tregs and RORγt+ Th17 cells. There were slight increases in FoxP3+RORγt-splenic Treg cells in mice on both the TL+Vehicle and TL+Indole diets compared to TS+Vehicle (Figure 6D and Supplemental Figure 8A); a ∼2-fold increase in the percentage of RORγt+FoxP3-splenic Th17 cells in the TL+Indole group compared to TL+Vehicle (Figure 6E and Supplemental Figure 8B); and ultimately leading to an increased Th17:Treg ratio (Figure 6F and Supplemental Figure 8C). These same trends were observed in colon T cells as well (Supplemental Figure 8D-F), suggesting a potential local (gut) and systemic (spleen) impact of Trp metabolism on Treg cells and indole-stimulated skewing of Th17 cells. The skewed splenic Th17:Treg ratio was only observed at CIA day 35, but not day 21 (Figure 6F), suggesting that the systemic Th17 response may be amplified following secondary immunization. Furthermore, re-stimulation of splenocytes harvested at day 35 with CII lead to significantly higher IL-17 production in the TL+Indole group compared to TL+Vehicle (Figure 6G), and a trend towards higher IL-17 in the TL+Indole group compared to TS+Vehicle. Although the trends were the same for stimulation with anti-CD3/CD28 beads (Figure 6H), the difference in IL-17 production between the groups was more pronounced with CII stimulation, suggesting a role for antigen-specific IL-17 production. There were no detectable differences in other Th17-related cytokines including IL-21 (below the limit of detection) and IL-22, although there was a trend towards increased GM-CSF in the TL+Indole group (Supplemental Figure 8G-L). Overall, these findings suggest that indole may either directly or indirectly skew the effector T cell response in CIA.

Our data thus far suggests that indole affects autoantibody sialylation, IgG2b production, and Th17 cellular responses. Because IL-23 has been shown to mediate Th17-derived IL-21 and IL-22 to enhance antibody pathogenicity via decreased sialylation of CII-specific antibodies [70], we hypothesized that indole mediates CIA induction through IL-23. To test this hypothesis, TL+Indole mice received anti-IL-23p19 or isotype control antibody weekly during pre-clinical CIA (days 0-21). As observed previously in CIA [83], anti-IL23p19 treatment resulted in a significant reduction in CIA severity compared to isotype control in TL+Indole mice (Figure 6I). Together, these data provide preliminary evidence that indole acts through the IL23/Th17 axis to incite disease via modulation of the autoantibody response.

### Indole stimulation induces similar pathways in human intestinal mononuclear cells

Finally, we tested whether indole could induce similar pathways in humans as well as mice. We rationalized that because indole is produced by the gut microbiome and absorbed through the gut, intestinal lymphocytes are more likely to interact with indole compared to circulating lymphocytes, and that testing the effects of indole on human colon mononuclear cells would be most physiologically relevant. We obtained lamina propria mononuclear cells (LPMCs) from macroscopically normal, discarded colon tissues obtained during bowel resection surgeries [84-86]. LPMCs from five donors were individually stimulated with 1mM indole or vehicle. After 4 hours, CD3+ T cells and CD19+ B cells were flow sorted from the pooled stimulation, RNA was extracted, sequenced, and Ingenuity Pathway Analysis was performed to identify differentially expressed pathways following indole stimulation compared to vehicle.

In the CD19+ B cell subset (Figure 7A and Supplemental Table 1), there were 21 pathways that were significantly enriched (p<0.05) and that were predicted to be significantly activated or repressed (z-score > |2|). Of these pathways, “IL-17 signaling” had the highest z-score (z=4.36, p=0.02), while “UDP-N-acetyl-D-glucosamine Biosynthesis II”, which produces the glycosylation precursor N-acetylglucosamine (GlcNAc), was the most statistically significant (z=2, p=0.0002). “NRF2-mediated Oxidative Stress Response”, “HIF1α Signaling”, “Unfolded Protein Response”, and “p38 MAPK Signaling” were among the other most significantly affected pathways . In the CD3+ T cell subset (Figure 7B and Supplemental Table 2), there were 3 pathways that significantly enriched and activated: “IL-17 signaling” (z=3.87, p=0.018), “Differential Regulation of Cytokine Production in Intestinal Epithelial Cells by IL-17A and IL-17F” (z=2.24, -p=0.003), and “Neuroprotective Role of THOP1 in Alzheimer’s Disease” (z=2.7, p=0.006). “Neuroprotective Role of THOP1 in Alzheimer’s Disease” was primarily driven by increased expression of endonucleases and serine proteases (Supplemental Table 2). Together, these data provide proof of concept that the observed indole-mediated effects on Th17 immunity, antibody glycosylation, and altered class switch recombination in mice with CIA may also be relevant for human intestinal immunity.

**Figure 7.**
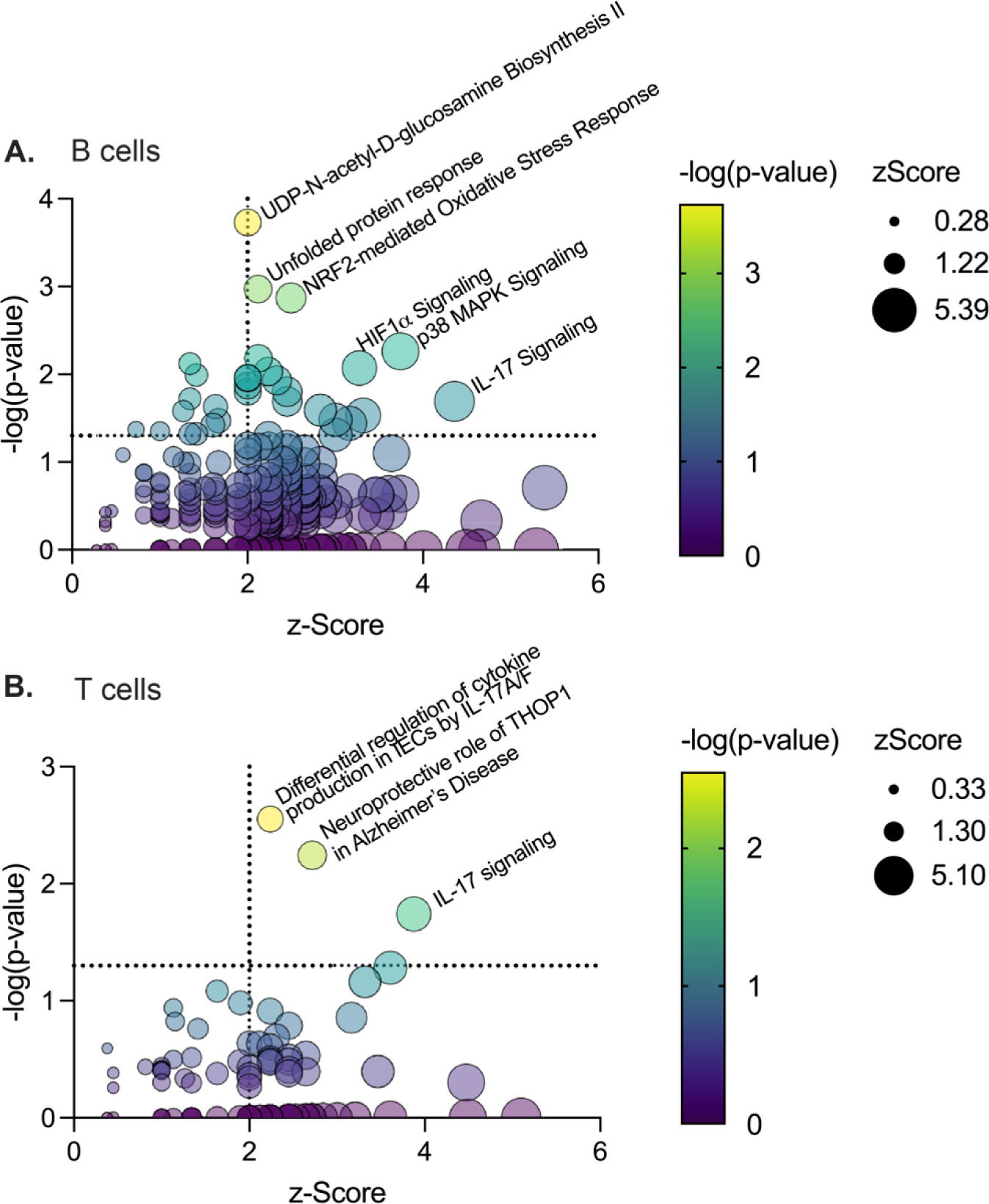
Indole-stimulated human intestinal cells also highlight changes in function. LPMCs isolated from healthy human colon tissue were stimulated with 1 mM indole or vehicle for 4hr. CD19+ B cells and CD3+ T cells were flow sorted, and RNA was isolated for RNAseq. Differentially expressed pathways (indole vs vehicle) were identified with Ingenuity Pathway Analysis for **(A)** CD19+ B cells and (**B)** CD3+ T cells. N=5 paired samples.

## Discussion

While there is compelling evidence for a mucosal origins hypothesis for RA and SpA, the mechanisms by which microbes contribute to the induction of autoimmunity remain elusive. There are several potential processes by which specific microbes could stimulate the development of autoimmunity, including altered production of bacterial metabolites, many of which have immunomodulatory functions [40-42]. Our observations of elevated Trp-derived indoles in two independent murine models of inflammatory arthritis (CIA and *Subdoligranulum isolate 7*-induced arthritis), as well as in patients with SpA [9], prompted us to investigate the requirement for indole in the development of CIA as a model in which cause-effect could be established. Because indole is exclusively produced from microbial metabolism of dietary Trp, we modulated indole production through either antibiotic-mediated microbiome depletion or reduction of dietary Trp. Depletion of either the microbiome or dietary Trp protected mice from developing CIA and adding back indole rescued CIA severity, suggesting that indole is sufficient to incite disease. These findings were confirmed further by amelioration of disease in mice colonized with Tryptophanase-deficient bacteria, which are unable to produce indole.

While altered Trp metabolism signatures have been observed in systemic lupus erythematosus (SLE), SpA, and RA [9, 37, 45, 47, 56-59], as well as in murine lupus and experimental autoimmune encephalitis (EAE) models [35, 48], there are conflicting roles for Trp metabolites in RA and experimental arthritis models. Indole-3-acetic acid (IAA), 5-hydroxytryptophan (5-HTP), and 5-HIAA ameliorate arthritis severity in proteoglycan-induced spondyloarthritis [49], CIA [50], and antigen-induced arthritis [44], respectively. Studies of kynurenine supplementation, or inhibition of kynurenine production with the IDO1 inhibitor 1-methyl-tryptophan (1-MT) have conflicting effects on arthritis severity: 1-MT treatment ameliorates disease in the K/BxN [51] murine model but exacerbates disease in the CIA model [52]. Intraarticular injection of indole, but not kynurenine, into rabbit knees induced RA-like synovitis [53]. Furthermore, these metabolites are either exclusively host-derived (5-HIAA), or can be produced by either host or microbiome (IAA, 5-HTP, kynurenine). As such, there is a great need to understand the specific role of tryptophan metabolism and microbiome-derived metabolites in the development of autoimmune arthritis.

Despite the conflicting role of Trp metabolites in arthritis models, our findings are in agreement with recent findings that a Trp-low diet protects against disease in murine models of EAE and lupus [35, 48]. While Trp can be metabolized by both host and microbial enzymes, our study provides novel evidence that microbial, rather than host, metabolism of Trp plays a major role in the development of disease in CIA, in agreement with what has previously been shown in models of SLE and EAE [48, 90, 91]. However, despite the consilience of these findings, it appears that the specific Trp metabolites that potentiate disease may be unique to each model. Because each of these disease models have been shown to be microbiome-dependent, the unique dysbiosis of each model may lead to differential production of Trp metabolites by the microbiota, and thus potentiate disease via distinct mechanisms: in lupus-prone triple congenic *B6.Sle1.Sle2.Sle3* (TC) mice, elevated levels of serum kynurenine appear to impair Treg function [35]. While kynurenine is thought to primarily be produced by host metabolism of Trp via indoleamine-2,3-dioxygenase (IDO1), the microbiota can also produce kynurenine via tryptophan-2,3-dioxygenase (TDO2) [92]. Choi et al. [35] show that antibiotic treatment, but not IDO1 inhibition, reduces serum kynurenine levels, suggesting that dysbiosis in TC mice leads to increased microbial-derived kynurenine that may promote disease development[35]. Alternatively, in an EAE model in which disease severity is exacerbated by colonization with *Lactobacillus reuteri*, Trp metabolism by microbial aromatic amino acid aminotransferase (ArAT) and aliphatic amidase E (AmiE) was increased in a Trp- and microbiome-dependent manner [48]. While both studies identify Trp metabolites that are correlated with disease severity and show that they can potentiate pathogenic T cell function in vitro, they do not test whether these metabolites directly incite disease in vivo. Thus, in our current study, we demonstrate for the first time the specific requirement for a Trp-derived metabolite in the development of CIA. Whether other indole-related metabolites can function similarly to indole in the CIA model, or if indole in the context of a complex metabolome will have similar effects will be essential to better understand the effects of Trp metabolites on host immunity both alone and in combination.

In our studies, we did not identify a specific indole-producing species that incites disease, but rather demonstrated a significant correlation between many taxa that were expanded in our CIA studies (*Firmicutes, Barnesiella, Rikenellaceae, Lactobacillales*) and indole levels. These taxa, as well as others that have been shown to be expanded in CIA (such as *Lachnospiraciae* and *Lactobacillaeae [32, 33]*), RA (*Lachnospiraceae bacterium, lactobacillus spp*., *Faecalibacterium*) [93, 94], and SpA (*Ruminococcaceae, Rikenellaceae, Porphyromonadaceae, Bacteroidaceae, Lachnospiraceae*) [4] are either predicted or known to produce indole (via TrpNet.ca), providing candidate bacteria for further study. Linking the essential indole-producing microbes to the development of inflammatory arthritis may provide a novel therapeutic opportunity.

CIA is a multi-faceted disease model. While CII-reactive autoantibodies that bind to cartilage and activate complement are the primary pathologic drivers of disease [68, 69], CII-reactive B cells, CD4+ T cells, and pro-inflammatory cytokines are also required for disease development. Indole supplementation appears to affect several aspects of CIA pathophysiology, including the induction of pro-inflammatory cytokines, CD4+ T cell skewing towards Th17 cells, and antibody-mediated complement activation. As the specific cellular target(s) of indole has not yet been identified, it is difficult to dissect which of these effects are directly initiated by indole stimulation, and which effects are secondary to an indole-induced pro-inflammatory environment. Furthermore, while our studies suggest that indole had minimal effects on microbiome composition, we cannot definitively rule out the possibility that indole supplementation altered the microbiome. Despite this limitation, the magnitude (effect size) of the microbiome changes observed with the TS vs TL diets were much higher compared to the differences between TL+Vehicle and TL+Indole, suggesting that indole supplementation had a much stronger effect on host immunity than it did on the microbiome.

Despite these confounding factors, our findings allow us to construct a hypothesized mechanism through which indole acts. We show increased complement deposition in the joints of indole-supplemented CIA mice, and increased complement activation by CII-specific antibodies. Our data suggest that indole enhances complement activation through two mechanisms: skewed CII-specific antibody isotype switching and glycosylation. First, indole restored CII-specific IgG2b production. Complement activation has been shown to differ by isotype, with IgG2b ≥ IgG2a > IgG1 [72-74]. Analysis of high-affinity anti-erythrocyte autoantibodies demonstrated that IgG2b antibodies had >200-fold increase in pathogenicity compared to other isotypes, which was almost entirely due to complement activation [77]. As such, the indole-mediated rescue of IgG2b production may explain the increase in complement fixation observed in anti-CII antibodies from the TL+Indole group.

Second, indole appears to alter antibody glycosylation. IgG-Fc sialylation acts as a protective “cap” to prevent complement binding to galactose residues. The ratio of galactosylated to sialylated residues was increased in CII-specific antibodies from indole supplemented mice, which may also explain the increase in complement fixation observed in anti-CII antibodies from the TL+Indole group. A prior study linked CII-antibody sialylation to IL-17 independent Th17 cellular responses during CIA: IL-23 stimulation of Th17 cells induced decreased IgG-Fc sialylation via decreased expression of *St6gal1* in B cells [70]. A similar study showed that IgG-Fc desialylation promotes nephropathy in an SLE mouse model in an IL-17A and IL-23-dependent manner [96]. We have previously shown that antibiotic-mediated protection from CIA is due, at least in part, to increased IgG-Fc sialylation, suggesting a role for microbiome-mediated changes in IgG-Fc sialylation [32]. As such, one possible explanation for our data is that microbiome-derived indole promotes IL23/Th17-mediated changes in antibody glycosylation. To support this explanation, we show that IL-23 blockade ameliorates disease in the indole-CIA model, suggesting that indole may act upstream of T cells, such as on IL-23-producing DCs. The observed expansion of Th17 cells in the indole supplemented group, and the requirement for IL-23 in indole-mediated CIA, provide further evidence for an effect of indole on the IL-23/Th17 axis in CIA. Furthermore, while the signals leading to IgG2b class switch recombination have not been well characterized in vivo, one study showed that IgG2b class switch recombination was decreased in IL-23p19-/- mice via decreased Tfh17 cells [97]. Altogether, these supporting data and ours suggest a mechanism in which indole induces pathogenic antibody formation through class switching to IgG2b as well as decreased antibody sialyation that enhances complement fixation, and that these changes may occur through the IL-23/Th17 axis. Whether (and how) Th17 cells directly affect antibody function in the indole-CIA model is yet to be determined, as prior studies have shown an IL-17-independent but IL-21- and IL-22-dependent role of Th17 cells on antibody glycosylation [70].

We also provide proof of concept that indole-mediated activation of immune cells may occur in humans as well. The pathways that were upregulated in human colon lymphocytes following indole stimulation paint a picture of Th17 cell and plasma cell activation, similar to what we observed in mice with CIA: “IL-17 signaling” (which was upregulated in both B and T cells) promotes B cell differentiation and plasma cell formation. “HIF1α signaling” and “NRF2-mediated oxidative stress response” were also upregulated in B cells: hypoxic environments induce HIF1α and NRF2, which then promote plasma cell formation [87]; the unfolded protein response (which was also upregulated in B cells) is upregulated during plasma cell differentiation to accommodate increased antibody production [98]. Furthermore, it has been shown that IL-17 signaling can sustain the plasma cell response via p38-MAPK signaling in B cells from lupus patients [88]; p38-MAPK signaling (upregulated in B cells) also regulates class switch recombination [89]. Finally, in concordance with our findings, individuals with RA have aberrant total IgG glycosylation that increases as individuals transition from the at-risk stage to disease and that correlates with disease activity [101], as well as an increased Th17 signature [24-26, 29, 30]. While further studies are required to validate which pathway(s)/genes are indeed upregulated following indole stimulation, and whether indole and primarily affects intestinal immunity (serving as a mucosal trigger) or systemic immunity (i.e. by exacerbating joint pathology), this data provides proof of concept that the increased Th17 signature and altered antibody patterns observed in CIA and indole-stimulated LPMCs may also be relevant to human disease.

Altogether, this study provides one potential mechanism by which the microbiome directly contributes to the development of autoimmunity and lays the groundwork for future mechanistic studies on the effect of indole on the IL23/Th17 axis and autoantibody pathogenicity, as well as the potential of blocking indole production as a therapy for RA and SpA.

## Methods

Detailed methods can be found in the Supplemental Materials. Sequencing data is deposited in the publicly accessible Sequencing Read Archive BioProject accession PRJNA1004672 (16S data) and the Gene Expression Omnibus accession GSE241655 (RNA sequencing).

**Statistics.** Unless specified otherwise, data was analyzed using GraphPad Prism software version 9; specific statistical tests for comparisons are referenced in the figure legends.

**Study Approval.** All animal studies were approved by the University of Colorado School of Medicine Institutional Animal Care and Use Committee (protocol #173). Research associated with the use of LPMC was reviewed by the Colorado Multiple Institutional Review Board (COMIRB protocol #17-0977) at the University of Colorado Anschutz Medical Campus and deemed Not Human Subject Research as defined by their polices in accordance with OHRP and FDA regulations.

## Author contributions

BJS, BT, WKJ, and KAK designed the project, performed the experiments, and analyzed the data. BJS and KAK assembled the data and wrote the manuscript. BA, JT, AS, SL, MEC, and SF contributed to data acquisition and analysis. AO, EEA, and SPC ran and analyzed the metabolites studies. DNF, JMK, and CER ran and analyzed the microbiome studies. ASD and SPC provided the *E. coli* mutants. SLS and RMA ran and analyzed the glycosylation studies. AJB, AS, SMD, and CCW procured, processed, and acquired and analyzed data for the human LPMC studies. All authors reviewed the manuscript and approved its final, submitted version.

## Acknowledgements

The work by the authors is supported through NIAMS R01AR075933 (KAK), NIAMS P30 AR079369, NIAMS T32AR007534 (WKJ, BT, AJB, MEC, SF), NIAID T32AI007405 (BJS), NIAID F30AI174817 (BJS), and the Rheumatology Research Foundation Future Physician Scientist Award (BJS, MEC) and Scientist Development Award (AJB).

