## Supplemental for "Microbiota-dependent indole production is required for the development of collagen-induced arthritis"

**Supplemental Methods**

**Animal studies.** *Collagen-induced arthritis.* Six-week-old male DBA/1 mice (The Jackson Laboratory or Envigo) were injected intradermally with 200µg bovine CII (Elastin) emulsified in 50µl complete Freund's adjuvant (CFA, Sigma) on days 0 and 21. Disease severity was measured as the sum of the clinical scores for each of the animal's four paws, where 0 = normal, 1 = one swollen digit, 2 = two swollen digits, 3 = three swollen digits, and 4 = entire paw swollen with ankylosis. Mice were euthanized at day 14, 21, or at the plateau of CIA severity (day 35-50) and feces, serum, and tissues were collected.

*Antibiotic treatment.* Antibiotics were administered as previously described[1]. Briefly, ampicillin (RPI, 1g/L) neomycin (RPI, 1g/L), vancomycin (Alfa Aesar, 0.5g/L), metronidazole (RPI, 0.5g/L), and grape-flavored Kool-Aid (20g/L) (Kraft Foods, to encourage consumption) were given in drinking water to mice beginning on day 21 through the end of the study. Kool-Aid alone was provided to the control group.

*Dietary intervention.* On CIA day -1, mice were given either Amino Acid chow (AA; Envigo, TD.01084, 0.18% L-Trp) or Trp Deficient Diet VI (Envigo, TD.130674). "Trp-Sufficient" (TS) mice were maintained on the AA diet for the duration of the experiment. "Trp-Low" (TL) mice were alternated between 5 days of Trp Deficient Diet VI and 2 days of Amino Acid diet for a cumulative 0.05% Trp-low diet. Indole (Acros Organics, 0.1mg/ml) was given in drinking water to mice beginning on day 21 through the end of the study. Alternatively, 200µl of 10mM indole in water was administered by oral gavage every other day beginning on day 0.

E. coli *colonization.* 6-week old germ-free DBA/1 mice were maintained on standard rodent show (Envigo 2020SX) containing 0.2% L-Trp. On day -7, mice received $10^8$ CFU *E. coli* BW25113 *ΔtnaA* or *E. coli* BW25113 *ΔbcsQ*. *ΔbcsQ* was selected as the isogenic control because *bcsQ* is a pseudogene in *E. coli* BW25113 and should have little to no phenotype in these studies. CIA was induced on day 0 after *E. coli* colonization was established.

**Microbiome Analysis.** Total genomic DNA was extracted using the QIAamp PowerFecal DNA kit (Qiagen Inc, Carlsbad, CA), which employs chemical and mechanical disruption (Roche MagNA Lyser) of biomass. PCR amplicons were generated using barcoded[2] primers targeting the V3V4 variable region of the 16S rRNA gene (338F: 5'ACTCCTACGGGAGGCAGCAG and 806R: 5' GGACTACHVGGGTWTCTAAT)[3,4]. PCR products were normalized using a SequalPrep™ kit (Invitrogen, Carlsbad, CA) and then pooled. The amplicon pool was partially lyophilized to reduce its volume, purified and concentrated using a DNA Clean and Concentrator Kit (Zymo, Irvine, CA), and then quantified using a Qubit Fluorometer 2.0 (Invitrogen, Carlsbad, CA). The pool was diluted to 4nM and denatured with 0.2 N NaOH at room temperature. The denatured DNA was diluted to 15pM and spiked with 25% of the Illumina PhiX control DNA prior to loading the sequencer. Paired-end sequencing was performed on the Illumina MiSeq platform with versions v2.4 of the MiSeq Control Software and of MiSeq Reporter, using a 600 cycle version 3 reagent kit.

Paired-end sequences were sorted by sample via barcodes in the paired reads with a Python script.[5,6] The paired reads were assembled using phrap[7,8] and pairs that did not assemble were discarded. Assembled sequence ends were trimmed over a moving window of 5 nucleotides until average quality met or exceeded 20. Trimmed sequences with more than 1 ambiguity or shorter than 250 nt were discarded. Potential chimeras identified with Uchime (usearch6.0.203_i86linux32)[9] using the Schloss[10] Silva reference sequences were removed from subsequent analyses. Assembled sequences were aligned and classified with SINA (1.3.0-r23838)[11] using the 418,497 bacterial sequences in Silva 115NR99[12] as reference configured to yield the Silva taxonomy; taxonomic assignments used the lowest-common-ancestor approach with default SINA settings. Closed-reference, operational taxonomic units were produced by binning sequences with identical Silva/SINA LCA assignments. Taxa with >0.01% abundance in any sample and observed in >5% of the samples were included in further analyses. All samples had a Good's coverage index >99%, indicating excellent depth of sequencing coverage.

The software packages R (v3.6.3) [13] and Explicet (v2.10.5) [14] were used to analyze and visualize data. Alpha-diversity indices (i.e., Chao1, Shannon H, Shannon H/Hmax) were evaluated between groups by ANOVA. Differences in overall microbiota composition (i.e., beta-diversity) were assessed through permutational ANOVA (PERMANOVA) with the Aitchison dissimilarity index applied to sequence count data.[15,16] Principal coordinates analysis (PCoA) was carried out using Aitchison dissimilarities and the *wcmdscale* function in the *vegan* R package.[16] Individual taxa differing between treatment groups were identified using the ANOVA-like differential expression (ALDEx2) R package[17,18]. The distribution of taxa in each sequence library was estimated through

1000 Dirichlet Monte Carlo re-samplings of sequence count data. To account for the compositional nature of microbiome sequence data, datasets were then subjected to a center log-ratio transformation with all features used as the denominator. P-values were adjusted for multiple comparisons using the false discovery rate method.[19] Effect size plots are derived from the outputs of ALDEx2 and represent the median effect sizes, calculated as the median between-group difference in CLR values between groups divided by the largest within-group difference in CLR values.[17,18]

**Metabolomics.** Cecal tip (tissue and contents, 30-100 mg) were harvested at day 35, flash-frozen, and stored at -80°C. Metabolomic analyses were performed via one of three methods as follows:

*HPLC*: Metabolites were extracted as previously described[20] with minor variations. Briefly, cecal tissue and contents were reconstituted in 200µl of HPLC-grade 80% methanol, sonicated for 3 x 3 second pulses (BioLogics Inc., 150 V/T Ultrasonic Homogenizer, power output ~20%), and then centrifuged at 12,000g for 5 minutes. The supernatant was saved and the extraction was repeated for a total of three rounds, producing a total of 600µl of extract. Samples were filtered through 5kDa spin columns (Amicon) and metabolites analyzed by HPLC. Analyses were performed on an Agilent Technologies 1260 Infinity HPLC using a Sepax Br-C18 column (120 Å, 4.6 x 250 mm). Mobile phase A, HPLC-grade water pH 7.0; mobile phase B, HPLC-grade acetonitrile; column temperature at 30 °C and flow rate of 1 ml/min. Chromatographic separation of the metabolites was performed using a gradient of 10% to 90% B in 30 min followed by washing and equilibration periods at the end of each run. The indole derivatives were detected by absorption at 280 nm and their absorbance spectra and retention times were confirmed by co-injection with authentic standards. Area under the curve (AUC) was calculated for each metabolite and normalized to starting sample weight.

*HPLC-MS*: Indole derivatives were quantified in mouse cecal samples using reversed-phase high-performance liquid chromatography with electrochemical coulometric array detection (EC-HPLC; CoulArray, Thermo Scientific, Waltham, MA). Cecal samples were extracted in 80% methanol and protein precipitate was removed by centrifugation at 15,000 x g. Separation was achieved using an Acclaim Polar Advantage II C18 column (Thermo, Waltham, MA) at a flow rate of 1 ml/min on a gradient of 10% to 55% acetonitrile in 50 mM sodium phosphate buffer, pH=3, containing 0.42 mM octanesulphonic acid as an ion-pairing agent. Calibration curves were composed by performing linear regression analysis of the peak area versus the analyte concentration. The data were quantified using the peak area in comparison to standards.

*LC-MS/MS*: cecal metabolites were analyzed by LC-MS/MS as described previously[21].

**Detection of serum or supernatant cytokines.** A multianalyte ELISA (MesoScale Diagnostics) was used to measure the levels of TNF (lower limit of detection, LLOD, 1.3 pg/ml), IL-1β (LLOD 3.1 pg/ml), IL-6 (LLOD 4.8 pg/ml), IL-17A (LLOD 0.3 pg/ml), IL-10 (LLOD 3.8 pg/ml), IFNγ (LLOD 0.16 pg/ml), IL-21 (LLOD 6.5pg/ml), IL-22 (LLOD 1.2 pg/ml), IL-23 (LLOD 4.9 pg/ml), and GM-CSF (LLOD 0.16pg/ml) in either undiluted serum or supernatant, according to the manufacturer's protocol.

**Detection of CII-specific antibodies.** Type II collagen-specific antibodies were detected in mouse serum at CIA day 35 using previously published methods[22]. Briefly, 96-well plates (Nunc MaxiSorp) were coated overnight at 4C with 5µg/ml ELISA-grade bovine CII (Chondrex), washed 3x with PBS + 0.05% Tween-20, and blocked for 4hr at 4C with 1% BSA in PBS. Serum samples were added to the wells at a dilution of 1:40,000 and incubated overnight at 4C with rocking. The plates were then washed, and horseradish peroxidase (HRP)-conjugated goat anti-mouse IgG, IgG1, IgG2a, or IgG2b (vendor) were added to the wells at a dilution of 1:10,000 for 4 hours at 4C. Following washing, 1X TMB ELISA substrate solution (BD Biosciences) was added to the wells and the plates were developed at room temperature. The reaction was stopped with 2N $H_2SO_4$ and read at 450nm and 570nm wavelengths. Pooled serum from mice with severe CIA was used to generate a standard curve in which the top standard was diluted 1:1000 (1unit/ml), followed by 2-fold serial dilutions.

**C3 activation ELISA.** Complement binding to CII-specific antibodies was assessed using previously published methods[1]. Ninety-six–well plates were coated overnight at 4°C with 5 µg/ml ELISA-grade mouse CII. The ELISA plates were then washed three times with 0.1% BSA + 0.05% Tween-20 in 1X PBS and blocked with 1% BSA in 1× PBS for 4 hours at 4°C. Serum samples were diluted 1:10,000 in 1× PBS. Samples were added to wells and incubated overnight at 4°C. The next day, wells were washed 5 times with 1× PBS + 0.05% Tween-20, then incubated with 15% normal mouse serum diluted in Dulbecco's PBS + 0.9 mM CaCl2 + 0.5 mM MgCl2 for 30 minutes at 37°C. The plates were then washed 5 times with 1× PBS and incubated with HRP-conjugated goat

IgG to mouse C3 (Cappel/MP Biomedicals) in 1:2,500 dilution for 1 hour at room temperature with rocker shaking. The plates were washed 5 times with 1× PBS and developed with 1× TMB ELISA substrate solution for 10 minutes. The reaction was then stopped with H2SO4 and read at 450 nm.

**Glycosylation studies.** Serum total IgG was purified by using Pierce Protein G Agarose (ThermoFisher Scientific) following the manufacturer's instructions. CII-specific antibodies were purified from serum as previously described[23] by coupling bovine CII (Chondrex) to CNBr-activated Sepharose 4B beads. To concentrate the eluted total IgGs, 3-kd Ultra-0.5 ml centrifugal filter units (Amicon) were used. Total N-linked glycan was released from glycoproteins using PNGase F (New England Biolabs) according to the manufacturer's instructions. Deglycosylation reactions were carried out at 37°C overnight to ensure effective release of glycans. Glycans were purified from the reaction using GlykoClean G Cartridges (Prozyme), dried, and fluorescence labeled with 2-aminobenzamide (Sigma-Aldrich). Labeled glycans were cleaned with GlykoClean S-plus Cartridges (Prozyme), dried, and subjected to high-performance liquid chromatography analysis. Glycan samples were dissolved in 25% 100 mM ammonium formate (pH 4.5) and 75% acetonitrile then separated using an Agilent 1260 Infinity Quaternary LC system outfitted with a 2.1 × 150 mm AdvanceBio Glycan Mapping column with 2.7 μm superficially porous particles and a fluorescence detector. Resulting peaks were analyzed in OpenLAB software (Agilent) and assigned glycoforms by comparing peaks of commercially available human IgG N-linked glycan library.

**Flow analysis.** Tissues were harvested from mice with CIA at day 21 or 35, homogenized in RPMI media, and passed through a 70 micron cell strainer. Red blood cells were lysed using Red Blood Cell Lysis buffer (eBioscience), and resuspended in FACS buffer (5% FBS in PBS) for surface staining. For intracellular staining, cells were fixed and permeabilized using the FoxP3/Transcription Factor Staining Buffer Kit (Tonbo). All antibodies and clones used are listed in Supplemental Table 3. Flow cytometric analysis was performed at the Barbara Davis Center BioResource Service Center, and analysis was performed using FlowJov10 software.

**Splenocyte re-stimulation.** Splenocytes were harvested as described above and re-stimulated with UV-crosslinked bovine Type II Collagen at a final concentration of 500ug/ml for 72 hours. Alternatively, $5x10^5$

splenocytes were stimulated with 2µl pre-washed CD3/CD28 Dynabeads. Supernatants were stored at -20C and were analyzed by multiplex immunoassay (MSD) as described above.

**Histopathology.** Mouse paws were removed at mid-limb and fixed in 10% paraformaldehyde. The bones were decalcified in 10% formic acid for 1 week and then embedded in paraffin. Sections of 5µm were cut from paraffin embedded tissues and stained with H&E. Pathology was assessed in a blinded manner for inflammation, pannus formation, and bone erosion. Pannus scoring criteria: 0 = no areas affected; 0.5=Very minimal, marginal zone only, less than 1% of area at risk affected; 1=Minimal infiltration of pannus in cartilage and subchondral bone, marginal zones mainly. Approximately 1-10% of area at risk affected; 2=Mild infiltration with marginal zone destruction of hard tissue in affected joints, 11-25% of area at risk affected; 3=Moderate infiltration with moderate hard tissue destruction in affected joints, 26-50% of area at risk affected; 4=Marked infiltration with marked destruction of joint architecture, affecting most joints, 51-75% of area at risk affected; 5=Severe infiltration associated with total or near total destruction of joint architecture, affects all joints, greater than 75% of area at risk affected. Bone resorption scoring criteria: 0.5=Very minimal resorption affects only marginal zones; 1=Minimal approximately 1-10% of area at risk of subchondral bone affected; 2=Mild, more numerous areas of resorption, approximately 11-25% of total area at risk of subchondral bone affected; 3=Moderate, obvious resorption of subchondral bone resulting in approximately 26-50% of area at risk of subchondral bone affected; 4=Marked, very obvious resorption of subchondral bone resulting in approximately 51-75% of area at risk of subchondral bone affected; 5=Severe, distortion of entire joint due to destruction approximately 76-100% of area at risk of subchondral bone affected. Inflammation scoring criteria: 0=no inflammatory infiltrate; 1=mild cellular infiltrate into joint and synovium; 2=enhanced cellular infiltrates, increased cell density throughout the joints, some joints affected; 3=maximal inflammation, high cell density, all joints affected.

**Complement C3 Immunohistochemistry.** Paraffin-embedded tissue slides were assessed for C3 complement deposition in the joints as described previously[24]. Complement deposition in each of the four paws was scored by a blinded observer from 0 to 3 (0=no staining, 1=mild staining, 2=moderate staining, 3=intense staining), and the average score across the four paws was plotted.

**Collection and Isolation of Human LPMC.** Colon tissue samples (N=5) were procured from the Program for Individuals with an Elevated Risk of Spondyloarthritis (PIERS) Registry. Healthy tissue was obtained from patients undergoing bowel surgery and would otherwise be discarded. These patients had no existing rheumatic disease or a history of Inflammatory Bowel Disease, HIV-1 infection, current treatment with immunosuppressive drugs, or recent chemotherapy (within 8 weeks). All patients undergoing surgery consented to the use of discarded tissue for research purposes. Protected patient information was de-identified to the laboratory investigators.

Lamina propria mononuclear cells (LPMC) were isolated from tissue samples as previously detailed[25-27]. Briefly, tissue specimens were trimmed of muscle and fat and treated with DL-Dithiothreitol (DTT; 1.67mM; Sigma-Aldrich) to remove additional mucus. The epithelial layer was subsequently removed with 1mM EDTA (Sigma-Aldrich) and the remaining tissue treated with collagenase D (0.5mg/ml, Roche Diagnostics). All released LPMCs were cryopreserved and stored in liquid nitrogen.

Cryopreserved LPMCs were thawed and stimulated at 37C with 1mM indole for 4 hours. CD3+ T cells and CD19+ B cells were flow sorted using a FACSAriaIII cell sorter. Total RNA was extracted using an RNeasy kit (Qiagen) according to the manufacturer's protocol. Libraries were constructed as previously described using the Next Ultra II directional RNA library prep kit with rRNA depletion[28]. Bulk RNA sequencing was performed on an Illumina MiSeq platform at the University of Colorado Genomics core. RNA-Sequencing workflow was implemented through the Bioconductor differential expression pipeline[29,30]. Salmon[31] was used for transcript quantification with GC bias correction, using a human transcript reference index from GENCODE (release 38) and no decoy sequences[32]. Tximeta[33] was used for importing transcripts which were then analyzed for differential expression with DESeq2[34]. Ingenuity Pathway Analysis as used to identify differentially expressed pathways. Differentially expressed pathways were defined as those with a p-value <0.05 and a z-score >2.

### Supplemental Table 1.

**Dataset: LPMC CD19+ B cells**

**UDP-N-acetyl-D-glucosamine Biosynthesis II**
**Pathway hits: 4/6, z-score: 2, p-value: 1.88E-04**

| Symbol | Entrez Gene Name | Ensembl | Expr Intensity/RPKM/FPKM/Counts | Expr Log Ratio | Expr p-value | Expr FDR (q-value) | Expr Fold Change | Expected | Location | Type(s) |
|---|---|---|---|---|---|---|---|---|---|---|
| GFPT1 | glutamine--fructose-6-phosphate transaminase 1 | ENSG00000198380.13 | 644.189 | 0.502 | 0.33 | 1 | 1.416 | Up | Cytoplasm | enzyme |
| GNPNAT1 | glucosamine-phosphate N-acetyltransferase 1 | ENSG00000100522.10 | 63.444 | 0.717 | 0.319 | 1 | 1.644 | Up | Cytoplasm | enzyme |
| PGM3 | phosphoglucomutase 3 | ENSG00000013375.16 | 179.912 | 0.558 | 0.4 | 1 | 1.472 | Up | Cytoplasm | enzyme |
| UAP1 | UDP-N-acetylglucosamine pyrophosphorylase 1 | ENSG00000117143.13 | 284.406 | 0.539 | 0.327 | 1 | 1.452 | Up | Nucleus | enzyme |

**Unfolded Protein Response**
**Pathway hits: 14/90, z-score: 2.121, p-value: 1.07E-03**

| Symbol | Entrez Gene Name | Ensembl | Expr Intensity/RPKM/FPKM/Counts | Expr Log Ratio | Expr p-value | Expr FDR (q-value) | Expr Fold Change | Expected | Location | Type(s) |
|---|---|---|---|---|---|---|---|---|---|---|
| DNAJB1 | DnaJ heat shock protein family (Hsp40) member B1 | ENSG00000132002.9 | 4872.28 | 0.509 | 0.469 | 1 | 1.423 | | Nucleus | transcription regulator |
| DNAJB13 | DnaJ heat shock protein family (Hsp40) member B13 | ENSG00000187726.9 | 15.777 | 0.681 | 0.054 | 1 | 1.603 | | Cytoplasm | other |
| DNAJC1 | DnaJ heat shock protein family (Hsp40) member C1 | ENSG00000136770.11 | 223.347 | 0.714 | 0.347 | 1 | 1.641 | | Cytoplasm | other |
| DNAJC3 | DnaJ heat shock protein family (Hsp40) member C3 | ENSG00000102580.15 | 615.599 | 0.514 | 0.484 | 1 | 1.428 | Down | Cytoplasm | other |
| DNAJC6 | DnaJ heat shock protein family (Hsp40) member C6 | ENSG00000116675.16 | 65.824 | 0.549 | 0.099 | 1 | 1.463 | | Cytoplasm | other |
| DNAJC16 | DnaJ heat shock protein family (Hsp40) member C16 | ENSG00000116138.13 | 184.32 | 0.51 | 0.04 | 1 | 1.424 | | Cytoplasm | other |
| ERO1B | endoplasmic reticulum oxidoreductase 1 beta | ENSG00000086619.14 | 140.026 | 0.51 | 0.377 | 1 | 1.424 | Up | Cytoplasm | enzyme |
| HSPA5 | heat shock protein family A (Hsp70) member 5 | ENSG00000044574.9 | 5794.735 | 0.529 | 0.478 | 1 | 1.443 | Up | Cytoplasm | enzyme |
| HSPA6 | heat shock protein family A (Hsp70) member 6 | ENSG00000173110.8 | 1338.989 | 1.191 | 0.121 | 1 | 2.283 | Up | Nucleus | enzyme |
| HSPA1A/H | heat shock protein family A (Hsp70) member 1A | ENSG00000204388.7 | 11949.356 | 0.845 | 0.298 | 1 | 1.796 | Up | Cytoplasm | enzyme |
| MAPK8 | mitogen-activated protein kinase 8 | ENSG00000107643.16 | 221.591 | 0.606 | 0.157 | 1 | 1.522 | Up | Cytoplasm | kinase |
| OS9 | OS9 endoplasmic reticulum lectin | ENSG00000135506.16 | 795.047 | 0.55 | 0.285 | 1 | 1.464 | | Nucleus | other |
| SEL1L | SEL1L adaptor subunit of ERAD E3 ubiquitin ligase | ENSG00000071537.14 | 1055.271 | 0.748 | 0.264 | 1 | 1.68 | Up | Cytoplasm | other |
| XBP1 | X-box binding protein 1 | ENSG00000100219.16 | 1737.376 | 0.743 | 0.329 | 1 | 1.674 | Up | Nucleus | transcription regulator |

**NRF2-mediated oxidative stress response**
**Pathway hits: 27/237, z-score: 2.496, p-value: 1.35E-03**

| Symbol | Entrez Gene Name | Ensembl | Expr Intensity/RPKM/FPKM/Counts | Expr Log Ratio | Expr p-value | Expr FDR (q-value) | Expr Fold Change | Expected | Location | Type(s) |
|---|---|---|---|---|---|---|---|---|---|---|
| CBR1 | carbonyl reductase 1 | ENSG00000159228.13 | 38.027 | 0.616 | 0.185 | 1 | 1.533 | Up | Cytoplasm | enzyme |
| CYP2C8 | cytochrome P450 family 2 subfamily C member 8 | ENSG00000138115.15 | 13.843 | 0.618 | 0.12 | 1 | 1.535 | | Cytoplasm | enzyme |
| CYP3A43 | cytochrome P450 family 3 subfamily A member 43 | ENSG00000021461.17 | 19.533 | 0.607 | 0.195 | 1 | 1.523 | | Cytoplasm | enzyme |
| DNAJB1 | DnaJ heat shock protein family (Hsp40) member B1 | ENSG00000132002.9 | 4872.28 | 0.509 | 0.469 | 1 | 1.423 | | Nucleus | transcription regulator |
| DNAJB13 | DnaJ heat shock protein family (Hsp40) member B13 | ENSG00000187726.9 | 15.777 | 0.681 | 0.054 | 1 | 1.603 | | Cytoplasm | other |
| DNAJC1 | DnaJ heat shock protein family (Hsp40) member C1 | ENSG00000136770.11 | 223.347 | 0.714 | 0.347 | 1 | 1.641 | | Cytoplasm | other |
| DNAJC3 | DnaJ heat shock protein family (Hsp40) member C3 | ENSG00000102580.15 | 615.599 | 0.514 | 0.484 | 1 | 1.428 | | Cytoplasm | other |
| DNAJC6 | DnaJ heat shock protein family (Hsp40) member C6 | ENSG00000116675.16 | 65.824 | 0.549 | 0.099 | 1 | 1.463 | | Cytoplasm | other |
| DNAJC16 | DnaJ heat shock protein family (Hsp40) member C16 | ENSG00000116138.13 | 184.32 | 0.51 | 0.04 | 1 | 1.424 | | Cytoplasm | other |
| ENC1 | ectodermal-neural cortex 1 | ENSG00000171617.15 | 107.579 | 0.584 | 0.061 | 1 | 1.499 | | Nucleus | peptidase |
| FOSL1 | FOS like 1, AP-1 transcription factor subunit | ENSG00000175592.9 | 54.077 | 0.556 | 0.337 | 1 | 1.47 | Down | Nucleus | transcription regulator |
| GSTA5 | glutathione S-transferase alpha 5 | ENSG00000182793.12 | 1.728 | 0.588 | 0.512 | 1 | 1.503 | | Cytoplasm | enzyme |
| GSTM1 | glutathione S-transferase mu 1 | ENSG00000134184.13 | 9.224 | 0.659 | 0.579 | 1 | 1.579 | | Cytoplasm | enzyme |
| HERPUD1 | homocysteine inducible ER protein with ubiquitin like d | ENSG00000051108.15 | 2285.948 | 0.644 | 0.246 | 1 | 1.562 | Up | Cytoplasm | other |
| HMOX1 | heme oxygenase 1 | ENSG00000100292.18 | 90.956 | 1.109 | 0.031 | 1 | 2.157 | Up | Cytoplasm | enzyme |
| KEAP1 | kelch like ECH associated protein 1 | ENSG00000079999.14 | 133.896 | 0.539 | 0.199 | 1 | 1.453 | Down | Cytoplasm | other |
| MAP2K3 | mitogen-activated protein kinase kinase 3 | ENSG00000034152.19 | 696.901 | 0.677 | 0.168 | 1 | 1.599 | Up | Cytoplasm | kinase |
| MAPK8 | mitogen-activated protein kinase 8 | ENSG00000107643.16 | 221.591 | 0.606 | 0.157 | 1 | 1.522 | Up | Cytoplasm | kinase |
| PIK3CG | phosphatidylinositol-4,5-bisphosphate 3-kinase catalyt | ENSG00000105851.11 | 770.839 | 0.868 | 0.144 | 1 | 1.825 | | Cytoplasm | kinase |
| PRKD1 | protein kinase D1 | ENSG00000184304.16 | 22.264 | 0.592 | 0.096 | 1 | 1.508 | Up | Cytoplasm | kinase |
| RASD1 | ras related dexamethasone induced 1 | ENSG00000108551.5 | 63.158 | 1.337 | 0.094 | 1 | 2.526 | | Cytoplasm | enzyme |
| RRAS | RAS related | ENSG00000126458.4 | 8.697 | 0.598 | 0.249 | 1 | 1.514 | Up | Cytoplasm | enzyme |
| SOD3 | superoxide dismutase 3 | ENSG00000109610.6 | 7.05 | 0.527 | 0.26 | 1 | 1.441 | | Extracellular | enzyme |
| SQSTM1 | sequestosome 1 | ENSG00000161011.20 | 3753.156 | 0.523 | 0.328 | 1 | 1.437 | Up | Cytoplasm | transcription regulator |
| TXNRD1 | thioredoxin reductase 1 | ENSG00000198431.16 | 596.526 | 0.923 | 0.168 | 1 | 1.897 | Up | Cytoplasm | enzyme |
| UBE2K | ubiquitin conjugating enzyme E2 K | ENSG00000078140.14 | 165.764 | 0.552 | 0.245 | 1 | 1.466 | Up | Cytoplasm | transcription regulator |
| USP14 | ubiquitin specific peptidase 14 | ENSG00000101557.15 | 262.135 | 0.544 | 0.309 | 1 | 1.458 | Up | Cytoplasm | peptidase |

**p38 MAPK signaling**
**Pathway hits: 15/118, z-score: 3.742, p-value: 5.47E-03**

| Symbol | Entrez Gene Name | Ensembl | Expr Intensity/RPKM/FPKM/Counts | Expr Log Ratio | Expr p-value | Expr FDR (q-value) | Expr Fold Change | Expected | Location | Type(s) |
|---|---|---|---|---|---|---|---|---|---|---|
| ATF1 | activating transcription factor 1 | ENSG00000123268.9 | 87.508 | 0.568 | 0.319 | 1 | 1.482 | Up | Nucleus | transcription regulator |
| BORCS8-M | BORCS8-MEF2B readthrough | ENSG00000064489.23 | 13.878 | 0.896 | 0.558 | 1 | 1.861 | Up | Nucleus | transcription regulator |
| FADD | Fas associated via death domain | ENSG00000168040.5 | 99.837 | 0.553 | 0.168 | 1 | 1.467 | | Cytoplasm | other |
| H3-4 | H3.4 histone, cluster member | ENSG00000168148.4 | 2.848 | 0.555 | 0.558 | 1 | 1.469 | Up | Nucleus | other |
| H3-3A/H3- | H3.3 histone A | ENSG00000132475.10 | 2802.453 | 0.724 | 0.162 | 1 | 1.652 | Up | Nucleus | other |
| HMGN1 | high mobility group nucleosome binding domain 1 | ENSG00000205581.11 | 574.47 | 0.561 | 0.258 | 1 | 1.475 | Up | Nucleus | transcription regulator |
| IL37 | interleukin 37 | ENSG00000125571.10 | 1.874 | 0.5 | 0.617 | 1 | 1.414 | Up | Extracellular | cytokine |
| IRAK4 | interleukin 1 receptor associated kinase 4 | ENSG00000198001.14 | 110.971 | 0.687 | 0.014 | 1 | 1.61 | Up | Cytoplasm | kinase |
| MAP2K3 | mitogen-activated protein kinase kinase 3 | ENSG00000034152.19 | 696.901 | 0.677 | 0.168 | 1 | 1.599 | Up | Cytoplasm | kinase |
| MEF2B | myocyte enhancer factor 2B | ENSG00000213999.17 | 38.523 | 1.149 | 0.027 | 1 | 2.218 | Up | Nucleus | transcription regulator |
| MEF2D | myocyte enhancer factor 2D | ENSG00000116604.18 | 1427.047 | 0.513 | 0.249 | 1 | 1.427 | Up | Nucleus | transcription regulator |
| PLA2G4B | phospholipase A2 group IVB | ENSG00000243708.11 | 24.702 | 0.909 | 0.064 | 1 | 1.877 | Up | Cytoplasm | enzyme |
| RPS6KA4 | ribosomal protein S6 kinase A4 | ENSG00000162302.13 | 101.57 | 0.537 | 0.189 | 1 | 1.451 | Up | Cytoplasm | kinase |
| RPS6KA5 | ribosomal protein S6 kinase A5 | ENSG00000100784.12 | 795.016 | 0.527 | 0.003 | 1 | 1.44 | Up | Nucleus | kinase |
| TAB2 | TGF-beta activated kinase 1 (MAP3K7) binding protein | ENSG00000228408.6 | 38.394 | 0.511 | 0.342 | 1 | 1.425 | Up | Cytoplasm | other |

**p53 signaling**
**Pathway hits: 13/98, z-score: 2.121, p-value: 6.55E-03**

| Symbol | Entrez Gene Name | Ensembl | Expr Intensity/RPKM/FPKM/Counts | Expr Log Ratio | Expr p-value | Expr FDR (q-value) | Expr Fold Change | Expected | Location | Type(s) |
|---|---|---|---|---|---|---|---|---|---|---|
| CASP6 | caspase 6 | ENSG00000138794.10 | 28.708 | 0.581 | 0.096 | 1 | 1.496 | Up | Cytoplasm | peptidase |
| CDKN1A | cyclin dependent kinase inhibitor 1A | ENSG00000124762.14 | 1191.713 | 0.598 | 0.266 | 1 | 1.514 | Up | Nucleus | kinase |
| GADD45A | growth arrest and DNA damage inducible alpha | ENSG00000116717.13 | 682.285 | 0.944 | 0.31 | 1 | 1.924 | | Nucleus | other |
| GADD45B | growth arrest and DNA damage inducible beta | ENSG00000099860.9 | 1537.727 | 0.53 | 0.385 | 1 | 1.444 | | Cytoplasm | other |
| GML | glycosylphosphatidylinositol anchored molecule like | ENSG00000104499.7 | 0.639 | 1.197 | 0.549 | 1 | 2.293 | Up | Plasma Mem | other |
| KAT2B | lysine acetyltransferase 2B | ENSG00000114166.8 | 303.612 | 0.774 | 0.127 | 1 | 1.71 | Up | Nucleus | transcription regulator |
| MAPK8 | mitogen-activated protein kinase 8 | ENSG00000107643.16 | 221.591 | 0.606 | 0.157 | 1 | 1.522 | Up | Cytoplasm | kinase |
| PCNA | proliferating cell nuclear antigen | ENSG00000132646.11 | 98.69 | 0.868 | 0.027 | 1 | 1.825 | | Nucleus | enzyme |
| PIK3CG | phosphatidylinositol-4,5-bisphosphate 3-kinase catalyt | ENSG00000105851.11 | 770.839 | 0.868 | 0.144 | 1 | 1.825 | Down | Cytoplasm | kinase |
| PPP1R13B | protein phosphatase 1 regulatory subunit 13B | ENSG00000088808.18 | 296.256 | 0.533 | 0.127 | 1 | 1.447 | | Cytoplasm | phosphatase |
| TIGAR | TP53 induced glycolysis regulatory phosphatase | ENSG00000078237.7 | 248.574 | 2.033 | 0.008 | 1 | 4.092 | | Cytoplasm | enzyme |
| TP73 | tumor protein p73 | ENSG00000078900.15 | 75.413 | 0.62 | 0.148 | 1 | 1.537 | Up | Nucleus | transcription regulator |
| TP53INP1 | tumor protein p53 inducible nuclear protein 1 | ENSG00000164938.14 | 742.269 | 0.586 | 0.269 | 1 | 1.501 | Up | Nucleus | other |

**HIF1α signaling**
**Pathway hits: 22/208, z-score: 3.273, p-value: 8.56E-03**

| Symbol | Entrez Gene Name | Ensembl | Expr Intensity/RPKM/FPKM/Counts | Expr Log Ratio | Expr p-value | Expr FDR (q-value) | Expr Fold Change | Expected | Location | Type(s) |
|---|---|---|---|---|---|---|---|---|---|---|
| ADRA1B | adrenoceptor alpha 1B | ENSG00000170214.5 | 15.618 | 0.749 | 0.136 | 1 | 1.681 | Up | Plasma Mem | G-protein coupled recep |
| BMP6 | bone morphogenetic protein 6 | ENSG00000153162.9 | 271.571 | 0.603 | 0.533 | 1 | 1.519 | Up | Extracellular | growth factor |
| CDKN1A | cyclin dependent kinase inhibitor 1A | ENSG00000124762.14 | 1191.713 | 0.598 | 0.266 | 1 | 1.514 | Up | Nucleus | kinase |
| HMOX1 | heme oxygenase 1 | ENSG00000100292.18 | 90.956 | 1.109 | 0.031 | 1 | 2.157 | Up | Cytoplasm | enzyme |
| HSPA5 | heat shock protein family A (Hsp70) member 5 | ENSG00000044574.9 | 5794.735 | 0.529 | 0.478 | 1 | 1.443 | Down | Cytoplasm | enzyme |
| HSPA6 | heat shock protein family A (Hsp70) member 6 | ENSG00000173110.8 | 1338.989 | 1.191 | 0.121 | 1 | 2.283 | Down | Nucleus | enzyme |
| HSPA1A/H | heat shock protein family A (Hsp70) member 1A | ENSG00000204388.7 | 11949.356 | 0.845 | 0.298 | 1 | 1.796 | Down | Cytoplasm | enzyme |
| IL6R | interleukin 6 receptor | ENSG00000160712.13 | 445.117 | 0.872 | 0.232 | 1 | 1.83 | Up | Plasma Mem | transmembrane recepto |
| LDHB | lactate dehydrogenase B | ENSG00000111716.14 | 188.01 | 0.548 | 0.224 | 1 | 1.462 | Up | Cytoplasm | enzyme |
| MAP2K3 | mitogen-activated protein kinase kinase 3 | ENSG00000034152.19 | 696.901 | 0.677 | 0.168 | 1 | 1.599 | Up | Cytoplasm | kinase |
| MMP1 | matrix metallopeptidase 1 | ENSG00000196611.6 | 15.476 | 0.559 | 0.069 | 1 | 1.473 | Up | Extracellular | peptidase |
| MMP10 | matrix metallopeptidase 10 | ENSG00000166670.10 | 1.989 | 1.157 | 0.167 | 1 | 2.229 | Up | Extracellular | peptidase |
| MMP15 | matrix metallopeptidase 15 | ENSG00000102996.5 | 37.677 | 0.876 | 0.009 | 1 | 1.836 | Up | Extracellular | peptidase |
| MMP25 | matrix metallopeptidase 25 | ENSG00000008516.18 | 18.581 | 0.616 | 0.22 | 1 | 1.532 | Up | Plasma Mem | peptidase |
| MMP28 | matrix metallopeptidase 28 | ENSG00000271447.6 | 17.636 | 0.64 | 0.084 | 1 | 1.558 | Up | Extracellular | peptidase |
| PIK3CG | phosphatidylinositol-4,5-bisphosphate 3-kinase catalyt | ENSG00000105851.11 | 770.839 | 0.868 | 0.144 | 1 | 1.825 | Up | Cytoplasm | kinase |
| PRKD1 | protein kinase D1 | ENSG00000184304.16 | 22.264 | 0.592 | 0.096 | 1 | 1.508 | Up | Cytoplasm | kinase |
| PROK1 | prokineticin 1 | ENSG00000143125.6 | 4.916 | 0.641 | 0.228 | 1 | 1.56 | Up | Extracellular | growth factor |
| RAC3 | Rac family small GTPase 3 | ENSG00000169750.9 | 2.296 | 0.764 | 0.432 | 1 | 1.698 | | Cytoplasm | enzyme |
| RASD1 | ras related dexamethasone induced 1 | ENSG00000108551.5 | 63.158 | 1.337 | 0.094 | 1 | 2.526 | Up | Cytoplasm | enzyme |
| RRAS | RAS related | ENSG00000126458.4 | 8.697 | 0.598 | 0.249 | 1 | 1.514 | Up | Cytoplasm | enzyme |
| SLC2A1 | solute carrier family 2 member 1 | ENSG00000117394.24 | 505.117 | 0.632 | 0.125 | 1 | 1.549 | Up | Plasma Mem | transporter |

| IL-17 signaling<br>Pathway hits: 19/187, z-score: 4.359, p-value: 2.03E-02 | | | | | | | | | | |
|---|---|---|---|---|---|---|---|---|---|---|
| **Symbol** | **Entrez Gene Name** | **Ensembl** | **Expr Intensity/RPKM/FPKM/Counts** | **Expr Log Ratio** | **Expr p-value** | **Expr FDR (q-value)** | **Expr Fold Change** | **Expected** | **Location** | **Type(s)** |
| CCL11 | C-C motif chemokine ligand 11 | ENSG00000172156.4 | 2.33 | 1.553 | 0.082 | 1 | 2.933 | Up | Extracellular | cytokine |
| CD40LG | CD40 ligand | ENSG00000102245.8 | 3.566 | 0.648 | 0.315 | 1 | 1.567 | Up | Extracellular | cytokine |
| CSF2 | colony stimulating factor 2 | ENSG00000164400.6 | 3.722 | 1.088 | 0.17 | 1 | 2.126 | Up | Extracellular | cytokine |
| DEFB119 | defensin beta 119 | ENSG00000180483.7 | 4.078 | 0.553 | 0.337 | 1 | 1.467 | Up | Extracellular | other |
| DEFB104A | defensin beta 104A | ENSG00000177023.2 | 1.792 | 0.762 | 0.433 | 1 | 1.695 | Up | Extracellular | other |
| DEFB107A | defensin beta 107A | ENSG00000186572.2 | 0.176 | 1.056 | 0.738 | 1 | 2.08 | Up | Extracellular | other |
| DEFB130A | defensin beta 130A | ENSG00000233050.1 | 0.401 | 1.041 | 0.556 | 1 | 2.057 | Up | Extracellular | other |
| DEFB4A/D | defensin beta 4A | ENSG00000171711.3 | 0.763 | 0.573 | 0.78 | 1 | 1.488 | Up | Extracellular | other |
| IL2 | interleukin 2 | ENSG00000109471.5 | 6.93 | 0.697 | 0.269 | 1 | 1.621 | Up | Extracellular | cytokine |
| IL3 | interleukin 3 | ENSG00000164399.5 | 2.863 | 0.817 | 0.24 | 1 | 1.762 | Up | Extracellular | cytokine |
| IL31 | interleukin 31 | ENSG00000204671.2 | 2.192 | 0.603 | 0.538 | 1 | 1.519 | Up | Extracellular | other |
| IL37 | interleukin 37 | ENSG00000125571.10 | 1.874 | 0.5 | 0.617 | 1 | 1.414 | Up | Extracellular | cytokine |
| MAP2K3 | mitogen-activated protein kinase kinase 3 | ENSG00000034152.19 | 696.901 | 0.677 | 0.168 | 1 | 1.599 | Up | Cytoplasm | kinase |
| MAPK8 | mitogen-activated protein kinase 8 | ENSG00000107643.16 | 221.591 | 0.606 | 0.157 | 1 | 1.522 | Up | Cytoplasm | kinase |
| PIK3CG | phosphatidylinositol-4,5-bisphosphate 3-kinase catalyt | ENSG00000105851.11 | 770.839 | 0.868 | 0.144 | 1 | 1.825 | Up | Cytoplasm | kinase |
| PROK1 | prokineticin 1 | ENSG00000143125.6 | 4.916 | 0.641 | 0.228 | 1 | 1.56 | Up | Extracellular | growth factor |
| RASD1 | ras related dexamethasone induced 1 | ENSG00000108551.5 | 63.158 | 1.337 | 0.094 | 1 | 2.526 | Up | Cytoplasm | enzyme |
| RRAS | RAS related | ENSG00000126458.4 | 8.697 | 0.598 | 0.249 | 1 | 1.514 | Up | Cytoplasm | enzyme |
| TAB2 | TGF-beta activated kinase 1 (MAP3K7) binding protein | ENSG00000228408.6 | 38.394 | 0.511 | 0.342 | 1 | 1.425 | Up | Cytoplasm | other |

#### Supplemental Table 2.

| Dataset: LPMC CD3+ T cells | | | | | | | | | | |
|---|---|---|---|---|---|---|---|---|---|---|
| **Differential regulation of cytokine production in intestinal epithelial cells by IL-17A/F** | | | | | | | | | | |
| **Pathway hits: 5/23, z-score: 2.236, p-value: 2.82E-03** | | | | | | | | | | |
| **Symbol** | **Entrez Gene Name** | **Ensembl** | **Expr Intensity/RPKM/FPKM/Counts** | **Expr Log Ratio** | **Expr p-value** | **Expr FDR (q-value)** | **Expr Fold Change** | **Expected** | **Location** | **Type(s)** |
| CCL4 | C-C motif chemokine ligand 4 | ENSG00000275302.2 | 100.812 | 0.656 | 0.303 | 1 | 1.575 | Up | Extracellular Space | cytokine |
| DEFB1 | defensin beta 1 | ENSG00000164825.4 | 1.345 | 2.582 | 0.031 | 1 | 5.988 | Up | Extracellular Space | other |
| DEFB4A/D | defensin beta 4A | ENSG00000177257.3 | 0.745 | 0.929 | 0.571 | 1 | 1.904 | Up | Extracellular Space | other |
| IFNG | interferon gamma | ENSG00000111537.5 | 50.736 | 0.651 | 0.399 | 1 | 1.57 | Up | Extracellular Space | cytokine |
| TNF | tumor necrosis factor | ENSG00000232810.4 | 131.681 | 0.519 | 0.286 | 1 | 1.433 | Up | Extracellular Space | cytokine |
| **Neuroprotective Role of THOP1 in Alzheimer's Disease** | | | | | | | | | | |
| **Pathway hits: 12/118, z-score: 2.714, p-value: 5.76E-03** | | | | | | | | | | |
| **Symbol** | **Entrez Gene Name** | **Ensembl** | **Expr Intensity/RPKM/FPKM/Counts** | **Expr Log Ratio** | **Expr p-value** | **Expr FDR (q-value)** | **Expr Fold Change** | **Expected** | **Location** | **Type(s)** |
| ENDOU | endonuclease, poly(U) specific | ENSG00000111405.9 | 26.739 | 0.858 | 0.045 | 1 | 1.813 | Up | Cytoplasm | peptidase |
| GNRH2 | gonadotropin releasing hormone 2 | ENSG00000125787.11 | 4.598 | 0.761 | 0.187 | 1 | 1.694 | Down | Extracellular Space | other |
| HGFAC | HGF activator | ENSG00000109758.9 | 3.314 | 0.69 | 0.321 | 1 | 1.613 | Up | Extracellular Space | peptidase |
| HPN | hepsin | ENSG00000105707.15 | 19.745 | 0.866 | 0.032 | 1 | 1.823 | Up | Plasma Membrane | peptidase |
| HTRA1 | HtrA serine peptidase 1 | ENSG00000166033.13 | 12.827 | 0.712 | 0.182 | 1 | 1.638 | Up | Extracellular Space | peptidase |
| IFNG | interferon gamma | ENSG00000111537.5 | 50.736 | 0.651 | 0.399 | 1 | 1.57 | Up | Extracellular Space | cytokine |
| KLK1 | kallikrein 1 | ENSG00000167748.11 | 22.834 | 0.553 | 0.115 | 1 | 1.467 | Up | Cytoplasm | peptidase |
| KLK8 | kallikrein related peptidase 8 | ENSG00000129455.15 | 12.375 | 0.508 | 0.164 | 1 | 1.422 | Up | Extracellular Space | peptidase |
| NTS | neurotensin | ENSG00000133636.11 | 3.627 | 0.973 | 0.164 | 1 | 1.963 | | Extracellular Space | other |
| PRSS57 | serine protease 57 | ENSG00000185198.12 | 3.812 | 0.588 | 0.479 | 1 | 1.504 | Up | Extracellular Space | peptidase |
| PRTN3 | proteinase 3 | ENSG00000196415.10 | 1.987 | 0.667 | 0.565 | 1 | 1.588 | Up | Extracellular Space | peptidase |
| TMPRSS9 | transmembrane serine protease 9 | ENSG00000178297.14 | 25.678 | 0.683 | 0.031 | 1 | 1.606 | Up | Plasma Membrane | peptidase |
| **IL-17 Signaling** | | | | | | | | | | |
| **Pathway hits: 15/187, z-score: 3.873, p-value: 1.83E-02** | | | | | | | | | | |
| **Symbol** | **Entrez Gene Name** | **Ensembl** | **Expr Intensity/RPKM/FPKM/Counts** | **Expr Log Ratio** | **Expr p-value** | **Expr FDR (q-value)** | **Expr Fold Change** | **Expected** | **Location** | **Type(s)** |
| CD40LG | CD40 ligand | ENSG00000102245.8 | 190.987 | 0.586 | 0.371 | 1 | 1.501 | Up | Extracellular Space | cytokine |
| DEFB1 | defensin beta 1 | ENSG00000164825.4 | 1.345 | 2.582 | 0.031 | 1 | 5.988 | Up | Extracellular Space | other |
| DEFB116 | defensin beta 116 | ENSG00000215545.1 | 0.849 | 1.125 | 0.489 | 1 | 2.182 | Up | Extracellular Space | other |
| DEFB124 | defensin beta 124 | ENSG00000180383.4 | 13.152 | 0.542 | 0.17 | 1 | 1.456 | Up | Extracellular Space | other |
| DEFB105A | defensin beta 105A | ENSG00000186562.8 | 7.298 | 0.56 | 0.248 | 1 | 1.474 | Up | Extracellular Space | other |
| DEFB107A | defensin beta 107A | ENSG00000198129.3 | 0.748 | 0.843 | 0.62 | 1 | 1.794 | Up | Extracellular Space | other |
| DEFB108B | defensin beta 108B | ENSG00000184276.3 | 2.1 | 1.188 | 0.166 | 1 | 2.279 | Up | Extracellular Space | other |
| DEFB4A/D | defensin beta 4A | ENSG00000177257.3 | 0.745 | 0.929 | 0.571 | 1 | 1.904 | Up | Extracellular Space | other |
| ERAS | ES cell expressed Ras | ENSG00000187682.2 | 2.042 | 0.784 | 0.515 | 1 | 1.722 | Up | Plasma Membrane | enzyme |
| IFNG | interferon gamma | ENSG00000111537.5 | 50.736 | 0.651 | 0.399 | 1 | 1.57 | Up | Extracellular Space | cytokine |
| MMP3 | matrix metallopeptidase 3 | ENSG00000149968.12 | 9.435 | 1.168 | 0.121 | 1 | 2.247 | Up | Extracellular Space | peptidase |
| OSM | oncostatin M | ENSG00000099985.4 | 16.33 | 0.563 | 0.196 | 1 | 1.477 | Up | Extracellular Space | cytokine |
| PIK3C2G | phosphatidylinositol-4-phosphate 3- | ENSG00000139144.11 | 28.442 | 0.517 | 0.205 | 1 | 1.43 | Up | Cytoplasm | kinase |
| TNF | tumor necrosis factor | ENSG00000232810.4 | 131.681 | 0.519 | 0.286 | 1 | 1.433 | Up | Extracellular Space | cytokine |
| TNFSF9 | TNF superfamily member 9 | ENSG00000125657.5 | 57.962 | 0.564 | 0.314 | 1 | 1.478 | Up | Plasma Membrane | cytokine |

**Supplemental Table 3**

| Antibody | Fluorophore | Clone | Vendor |
|---|---|---|---|
| CD4 | BUV395 | 30-F11 | BD Bioscience |
| PD-1 | BV421 | 29F.1A12 | Biolegend |
| Viability | Ghost | | Tonbo |
| CD44 | BV605 | IM7 | Biolegend |
| RORγt | BV786 | Q31-378 | BD Bioscience |
| CD25 | FITC | PC61.5 | Tonbo |
| CD3 | PerCPCy5.5 | 145-2C11 | Tonbo |
| CD69 | PE-Dazzle594 | H1.2F3 | Biolegend |
| FoxP3 | PE-Cy7 | 3G3 | Tonbo |
| Bcl6 | Alexa-Fluor 647 | IG191E/A8 | Biolegend |
| CD62L | APC-Cy7 | MEL-14 | Biolegend |

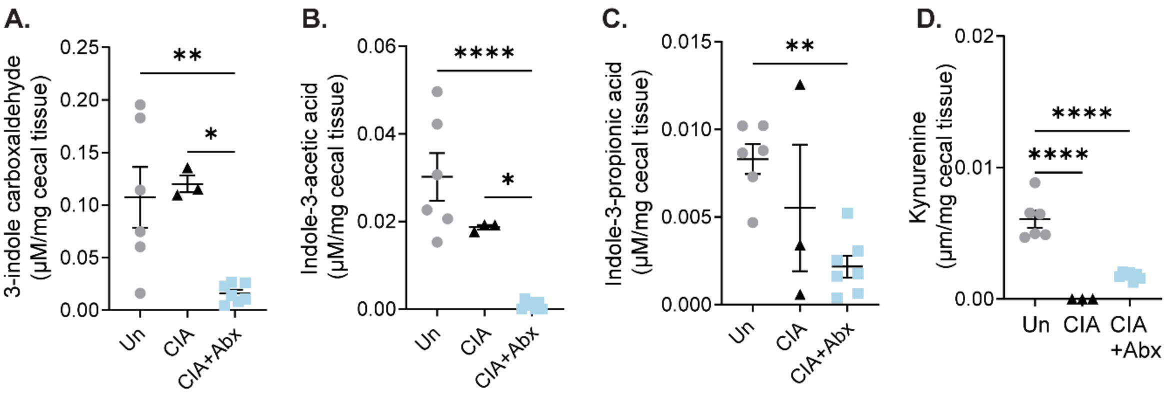


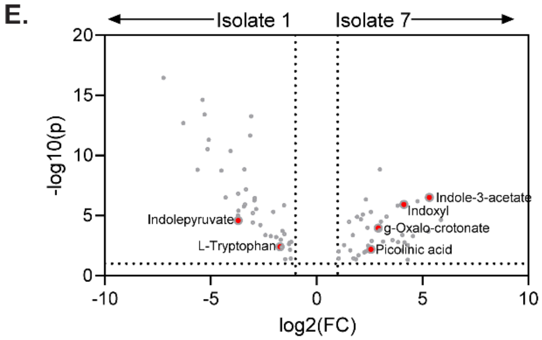


F.

| Phyla:Family | Spearman r | P-value |
|---|---|---|
| Firmicutes | 0.729994953 | 0.000583284 |
| Bact:Barnesiella | 0.647508383 | 0.003669957 |
| Bact:Rikenellaceae | 0.643883586 | 0.003930948 |
| Firm:Lactobacillales | 0.623292267 | 0.005717237 |
| Bact:VC2.1-Bac22 | 0.559292734 | 0.015812919 |
| Firm:Hydrogenoanaerobacterium | 0.508687437 | 0.031101164 |

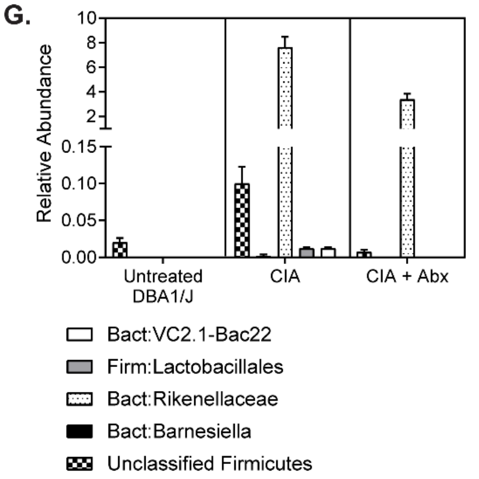


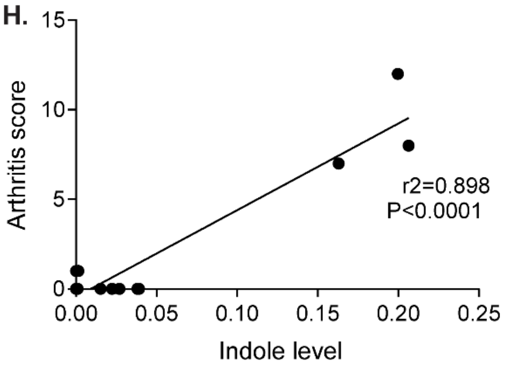


**Supplemental Figure 1. CIA induced microbiome-dependent changes in Trp metabolism.** CIA was induced in male 6-week old DBA/1J mice by injection of bovine type II collagen (CII) in complete Freund's adjuvant at days 0 and 21. Cecal contents were harvested at day 35 from CIA mice (n=3-5), CIA mice depleted from microbiota by antibiotic administration after day 21 (CIA+Abx, n=7), or untreated DBA/1J mice (Un, n=6). **(A-D)** HPLC was used to quantify Trp pathway metabolites indicated on the y-axis in μm. All data were reported as individual mice (symbols) and mean ±SEM (bars) after normalization to weight (mg) of cecal contents. *, p<0.05; ***, p<0.001; ****, p<0.0001 as determined one-way ANOVA with Tukey's multiple comparisons test. **(E)** Germ free DBA1 mice were orally gavaged at day 0 with either sterile PBS or $10^7$ CFU *Subdoligranulum didolesgii Isolate 1 or Isolate 7.* LC-MS/MS were used to screen >190 metabolites in cecal contents. Differential abundance of metabolites is shown as a volcano plot. **(F)** Paired 16S amplicon sequencing + LC-MS/MS metabolomics analysis were performed on mice with CIA compared to CIA + Abx. Spearman correlation revealed the top 6 OTUs that significantly correlated with levels of indoxyl. N=5-7 per group. **(G)** Relative abundance of the OTUs identified in (F). **(H)** Indole was measured by HPLC in the cecal contents of untreated DBA/J mice and mice with CIA ± Abx. Pearson r correlation of arthritis severity vs indole levels is shown.

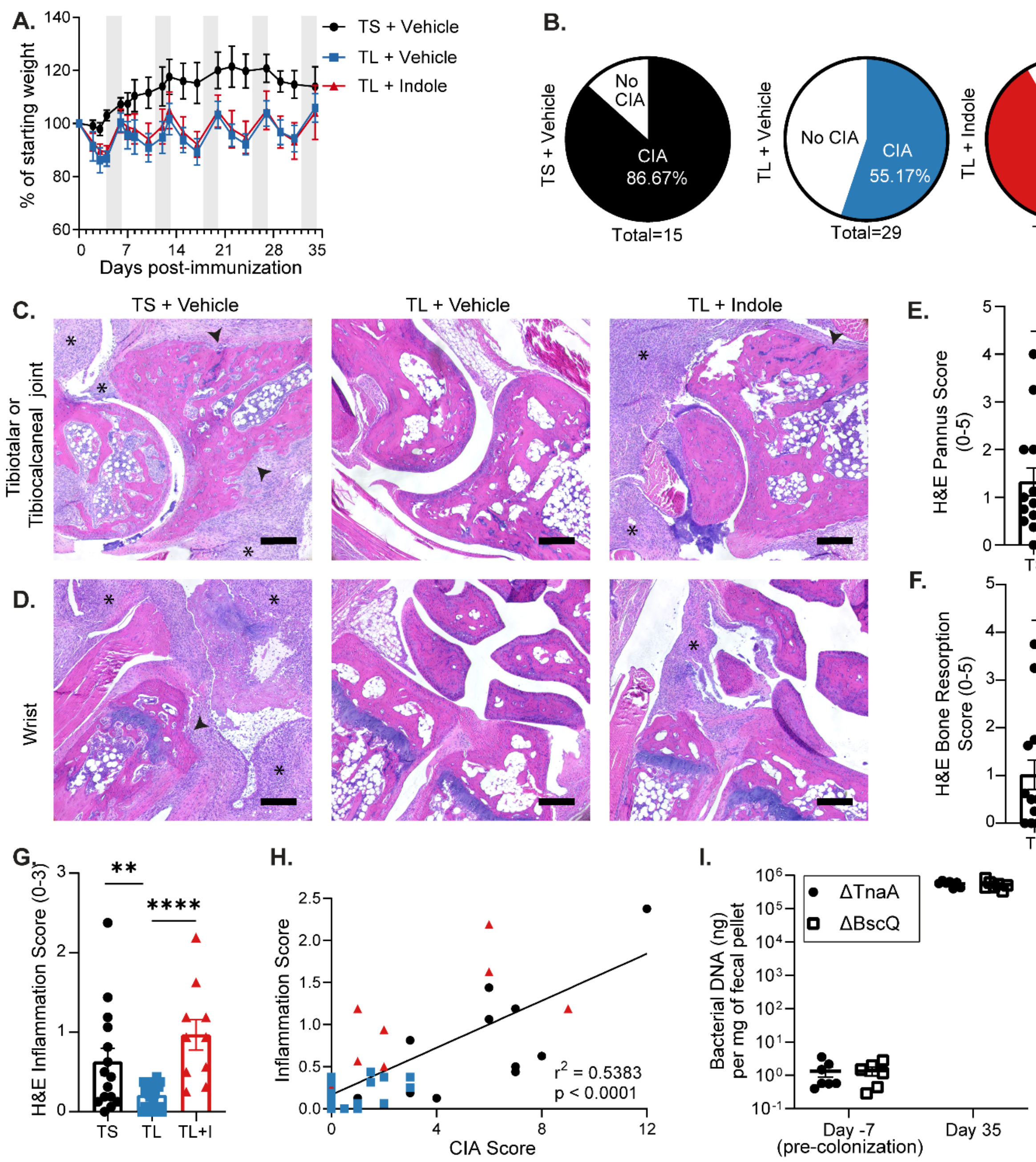


**Supplemental Figure 2. Indole is required for CIA.** Male 6-week old DBA/1J mice were fed a tryptophan-low (TL) diet or a trp-sufficient (TS) diet starting at day -1 through the duration of the experiment. CIA was induced by injection of CII in CFA at days 0 and 21. Indole (200µl of a 10mM solution) or vehicle control (0.33% methanol) was added back by oral gavage every other day starting on day 0. **(A)** % of starting body weight of mice with CIA on TL + vehicle, TL + indole, and TS + vehicle diets for the duration of the CIA study. In the TL treatment, after 5 days of Trp-deficient diet, mice are fed Trp-sufficient diet for 2 days (represented as grey bars) for a cumulative Trp-low diet. All values are plotted as mean ± SEM with n=10 (TL + vehicle), n=10 (TL + indole), n=5 (TS + vehicle) mice from one representative experiment. P<0.0001 by two-way ANOVA with Bonferroni adjustment for multiple comparisons for TL+Vehicle vs TS+Vehicle and TL+Indole vs TS+Vehicle. There was no statistical significance between TL+Vehicle and TL+Indole. **(B)** CIA incidence of mice in Figure 2B as defined by CIA score >1 at day 35, pooled from three independent experiments. **(C-D)** Representative H&E images of the tibiotalar joint **(C)** and wrist **(D)** for each group. Scale bar = 200µm; asterisks = synovial inflammation; arrowheads = bone resorption. **(E-G)** H&E stained paws were assessed for pannus formation (0-5), bone resorption (0-5), and inflammation (0-3), respectively. Each data point represents the average score of 4 paws per mouse. N=10-20 per group. **(H)** Correlation between CIA score (clinical) and inflammation score (H&E). **(I)** Colonization of *E. coli* BW25113 *ΔtnaA* or *E.coli* BW25113 *ΔbcsQ* in germ-free mice was verified by universal rpoB primers at CIA day 35 compared to pre-colonization (day-7). No significant differences were identified by unpaired t-test.

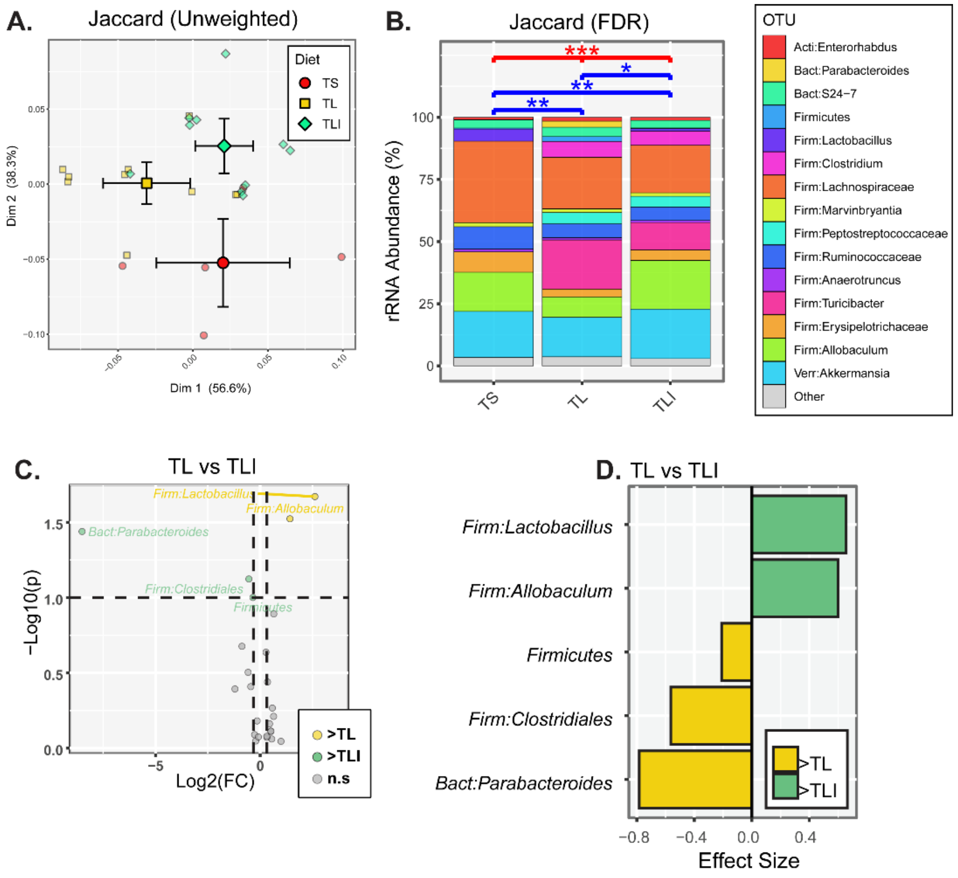


**Supplemental Figure 3. Indole minimally affects bacterial dysbiosis imparted by a TL diet during CIA.** The fecal microbiomes of mice in Figure 3 were also analyzed by Jaccard (unweighted) beta-diversity indices. **(A)** PCoA in which smaller, fainter symbols represent individual mice while larger symbols represent group means + 95% confidence intervals for PC1 and PC2. N=5-10 per group. **(B)** Bar charts annotated with results of PERMANOVA tests. *, p<0.05; **, p<0.01 and ***, p<0.001. **(C-D)** Volcano **(C)** and effect size **(D)** plots generated by ANOVA-like differential expression (ALDEx2) analysis indicate taxa that were enriched or depleted in TL + vehicle mice compared to TL + Indole. No taxa were found to be differentially abundant between TL + vehicle and TL + Indole groups after adjusting p-values for multiple comparisons (i.e., all FDR-corrected p values were >0.05), so p-value rather than FDR is shown in the volcano plot. N=5-10 per group.

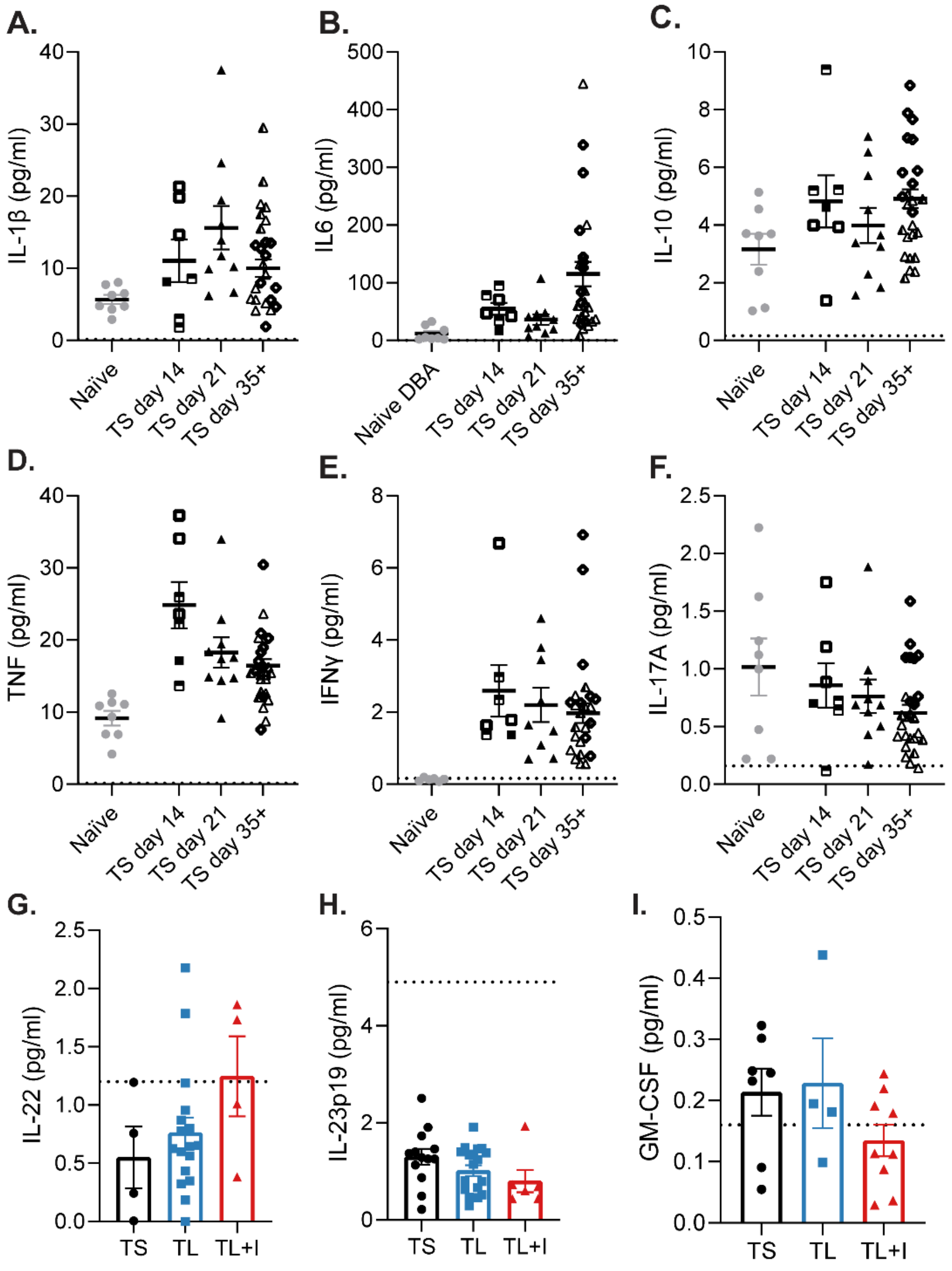


**Supplemental Figure 4. Indole alters the cytokine profile in CIA. (A-F)** Terminal serum was collected from male 6-week old DBA1 mice with CIA fed TS diet and treated with vehicle control (0.33% methanol) on CIA day 14, 21, and 35+. Naïve, age-matched, unimmunized male DBA1 mice were used as controls. Serum was analyzed by an 8-plex immunoassay (Mesoscale). The dashed line on the Y-axis denotes the lower limit of detection for each analyte. N=7-28 per timepoint plotted as individual mice (symbols) and mean ±SEM (bars). Each unique symbol represents an independent experiment. **(G-I)**, IL-22, IL-23p19, and GM-CSF were measured at day 35 only.

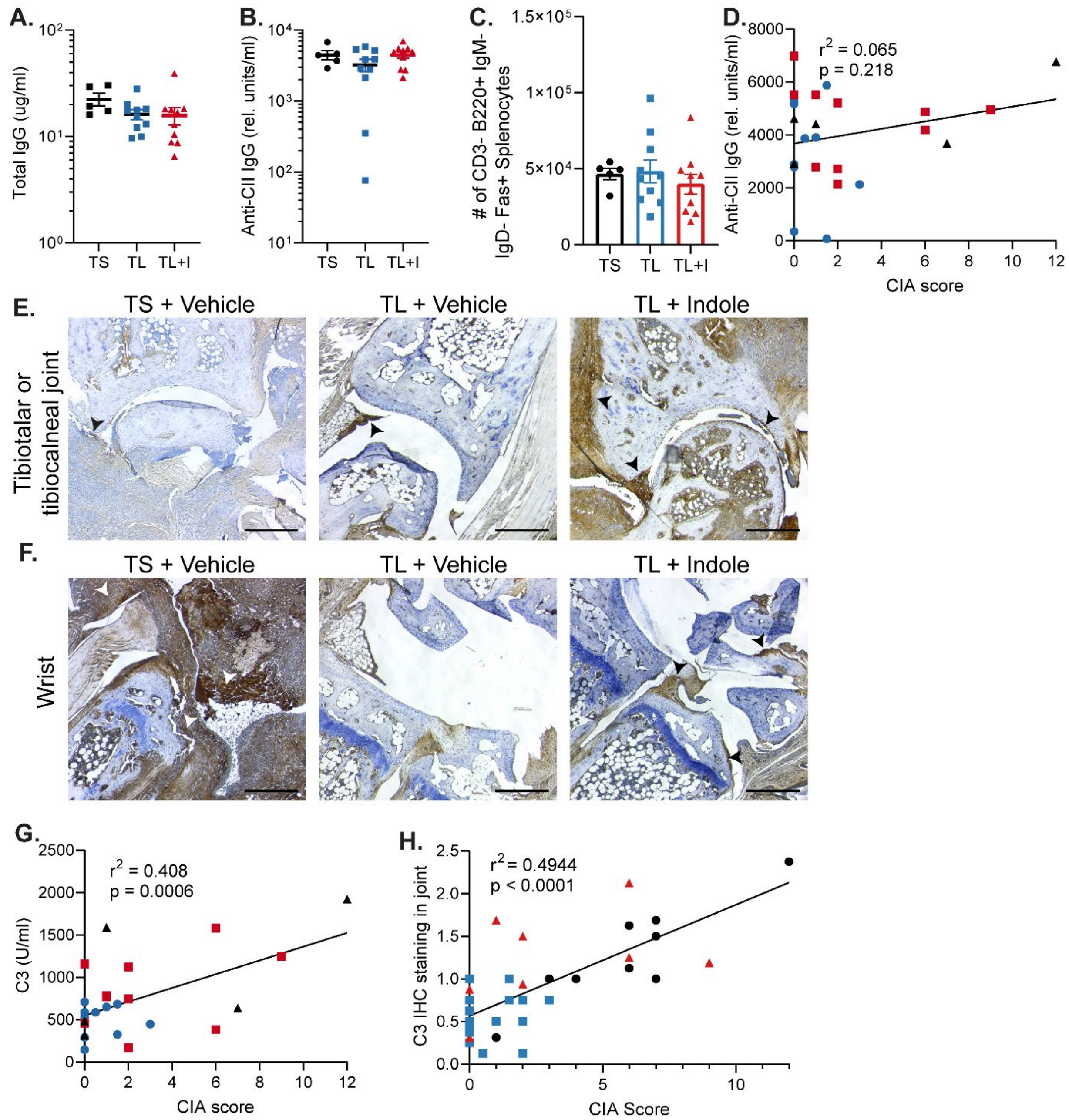


**Supplemental Figure 5. Antibody isotype and complement fixation correlate with development of CIA.** Day 35 serum from mice with CIA treated with TS + vehicle, TL + vehicle, and TL + indole was evaluated by ELISA for: **(A)** total IgG, **(B)** CII-specific IgG. **(C)** Splenocytes from mice with CIA fed a TL or TS diet and treated with indole or vehicle were counted and germinal center B cells (live CD3- B220+ IgM- IgD- Fas+) were assessed by flow cytometry. N=5-10 per group plotted as individual mice (symbols) and mean ±SEM (bars). No statistical significance was observed by unpaired t-test. **(D)** Anti-CII IgG levels were plotted against CIA score (x-axis). Simple linear regression was performed, and statistical p-value and $r^2$ value are shown. **(E-F)** Representative images of FFPE paws that were stained by immunohistochemistry for complement C3 (brown) and hematoxylin (blue). Scale bar = 200μm. Arrowheads = complement deposition. **(G)** CII-specific C3 activation was measured as described in Figure 5 and plotted against CIA severity. **(H)** C3 deposition by IHC was scored as described in figure 5 and plotted against CIA severity.

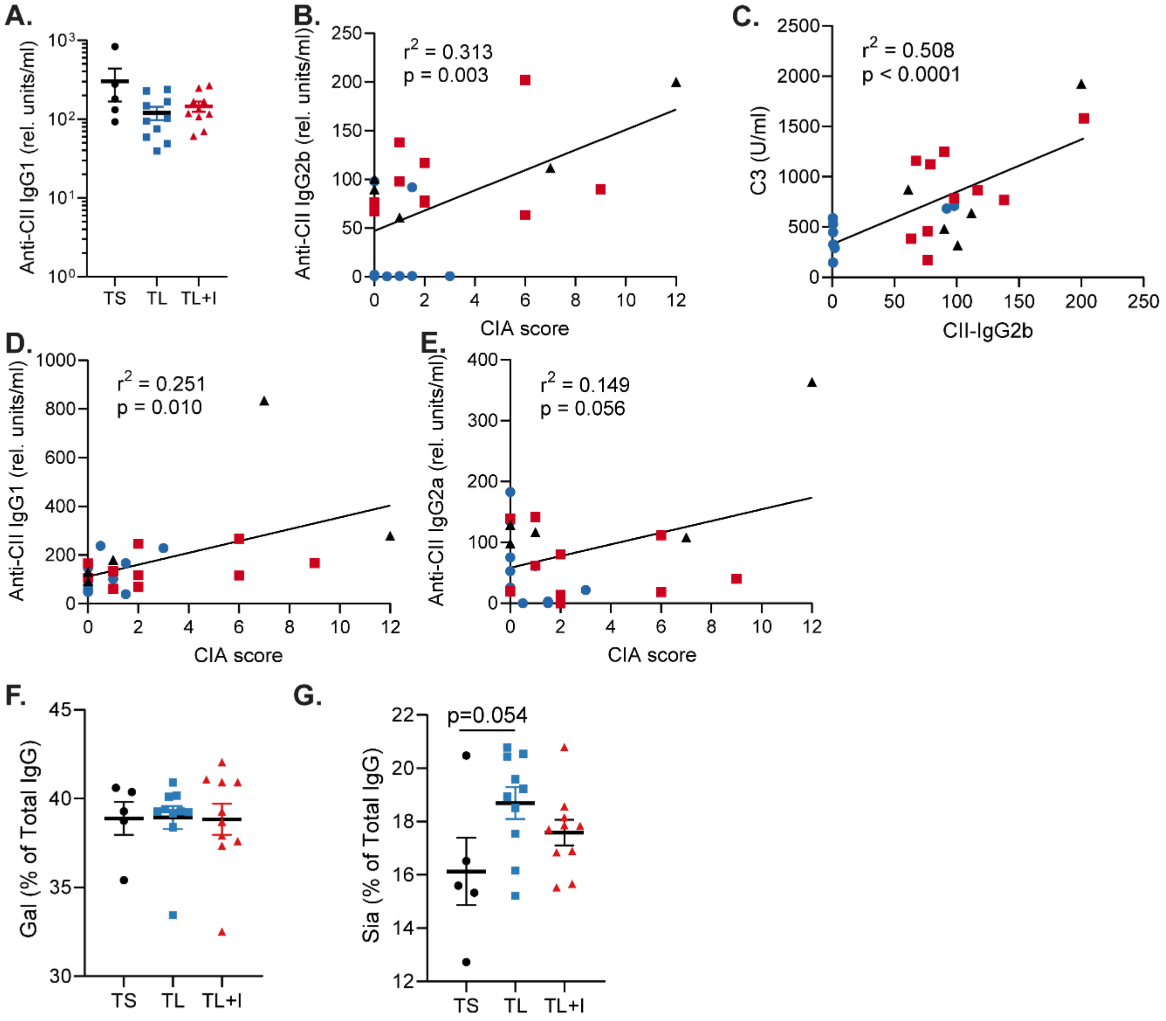


**Supplemental Figure 6. Indole alters complement activation by CII-specific antibodies as well as IgG subclass and glycosylation. (A)** Anti-CII IgG1 was measured in serum by ELISA at CIA day 35. **(B-C)** Anti-CII IgG2b was measured as described in Figure 5 and plotted against CIA score **(B)** and C3 activation **(C)**. Simple linear regression was performed, and statistical p-value and $r^2$ value are shown. **(D-E)** Anti-CII IgG1 and IgG2a were measured as described in Figure 5 and plotted against CIA score. Simple linear regression was performed, and statistical p-value and $r^2$ value are shown. **(F-G)** Total IgG was purified from serum and IgG glycosylation patterns were assessed by liquid chromatography with mass spectrometry (LC-MS/MS). Galactosylation and Sialylation were calculated as a % of all glycoforms (G0, G1, G2, S1, and S2). N=5-10 per group plotted as individual mice (symbols) and mean ±SEM (bars).

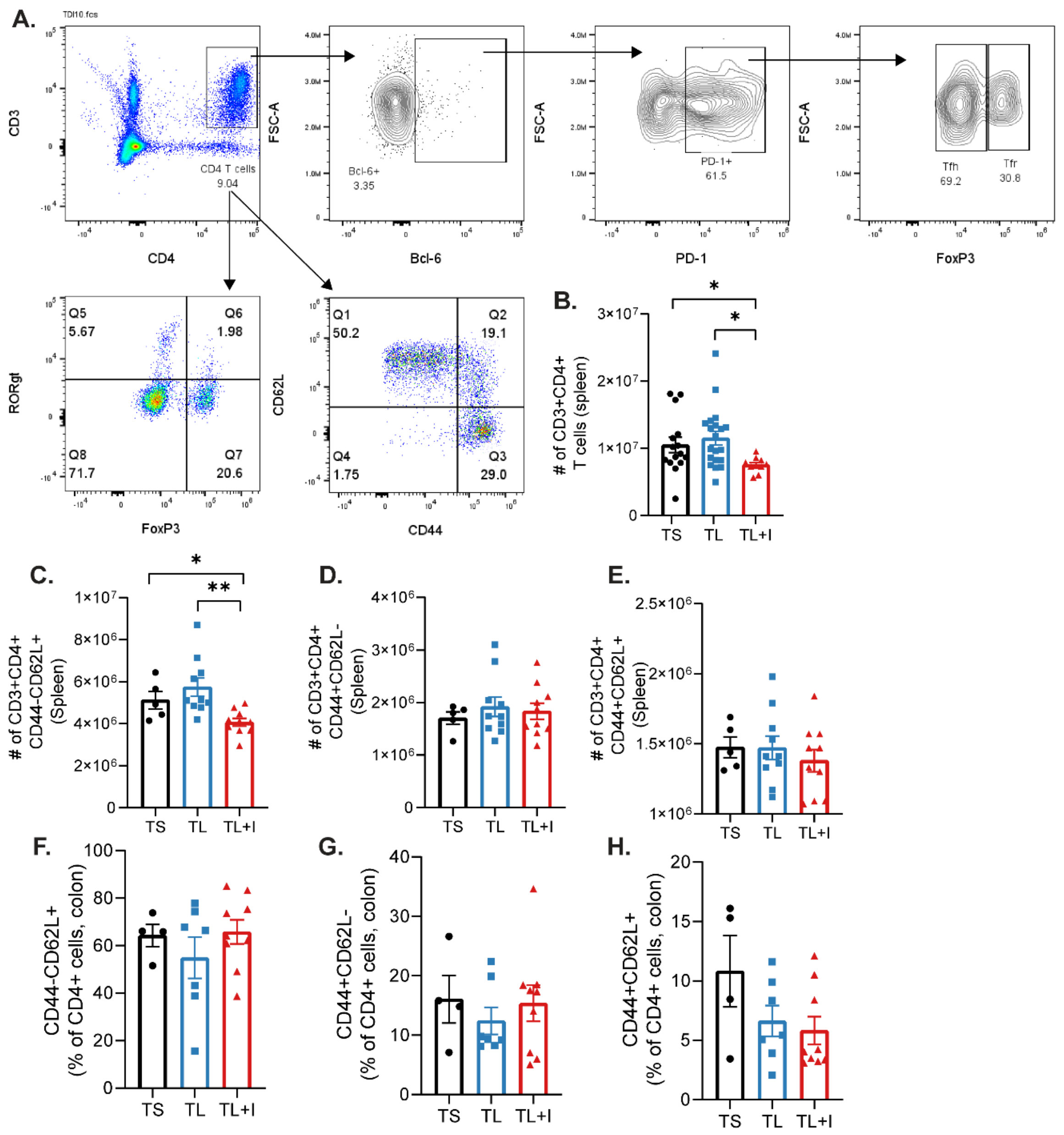


**Supplemental Figure 7. Indole skews effector T cell populations.** Splenocytes and colon LPMCs from mice with CIA fed a TL or TS diet and treated with indole or vehicle were analyzed by flow cytometry. **(A)** Representative gating strategy for Figure 6. **(B)** Total splenic CD3+CD4+ T cell counts at CIA day 35. N=10-20 per group. **(C)** Splenic $T_{\text{naïve}}$ (CD44-CD62L+) as # of CD4+ T cells. **(D)** Splenic $T_{\text{effector}}$ (CD44+CD62L-) as # of CD4+ T cells. **(E)** Splenic $T_{\text{CM}}$ (CD44+CD62L+) as # of CD4+ T cells. **(F)** Colon $T_{\text{naïve}}$ (CD44-CD62L+) as % of CD4+ T cells. **(G)** Colon $T_{\text{effector}}$ (CD44+CD62L-) as % of CD4+ T cells. **(H)** Colon $T_{\text{CM}}$ (CD44+CD62L+) as % of CD4+ T cells. N= 4-10 per group plotted as individual mice (symbols) and mean ± SEM (bars). *, $p<0.05$; **, $p<0.01$ by unpaired t-test.

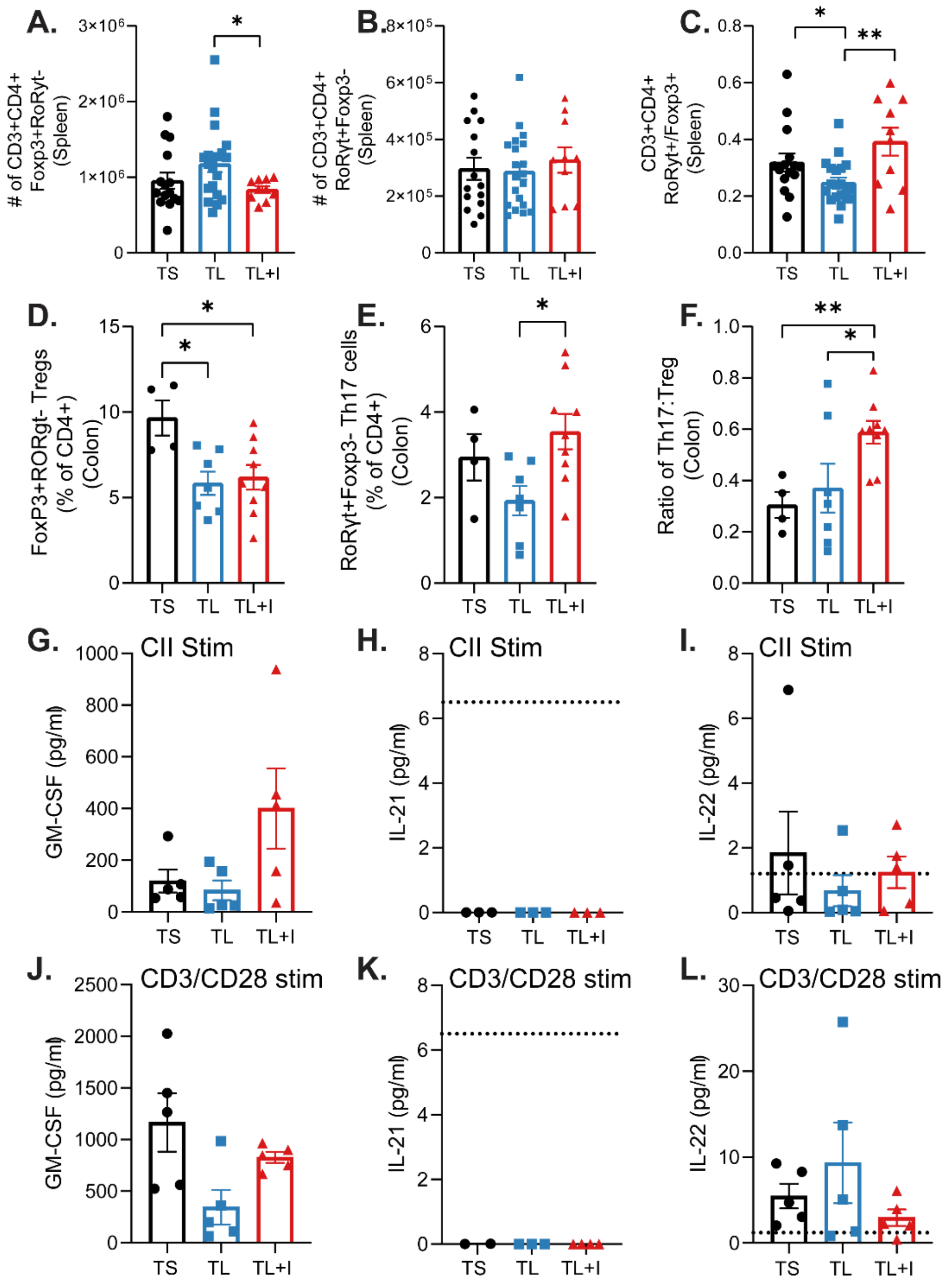


**Supplemental Figure 8. Indole skews towards Th17 cells. (A)** Total # of splenic CD3+CD4+FoxP3+RORγt-Treg cells **(B)** Total # of splenic FoxP3-RORγt+Th17 cells. **(C)** Ratio of the # of splenic Th17 cells:Treg cells. **(D)** Colon FoxP3+RORγt-CD25+ regulatory T cells are plotted as the percent of total CD4+ T cells. **(E)** Colon CD3+CD4+FoxP3-RORγt+ Th17 cells are plotted as the percent of total CD4+ T cells. **(F)** Ratio of colon Th17 to Treg cells. N=10-20 per group (spleen) and 4-10 per group (colon) plotted as individual mice (symbols) and mean ± SEM (bars). **(G-L)** Total splenocytes from CIA day 35 were harvested and re-stimulated with bovine type II collagen **(G-I)** or CD3/CD28 Dynabeads **(J-L)**; supernatant was saved and cytokines (as denoted on the y-axis) were measured by MSD**.** N=5 per group from one representative experiment.

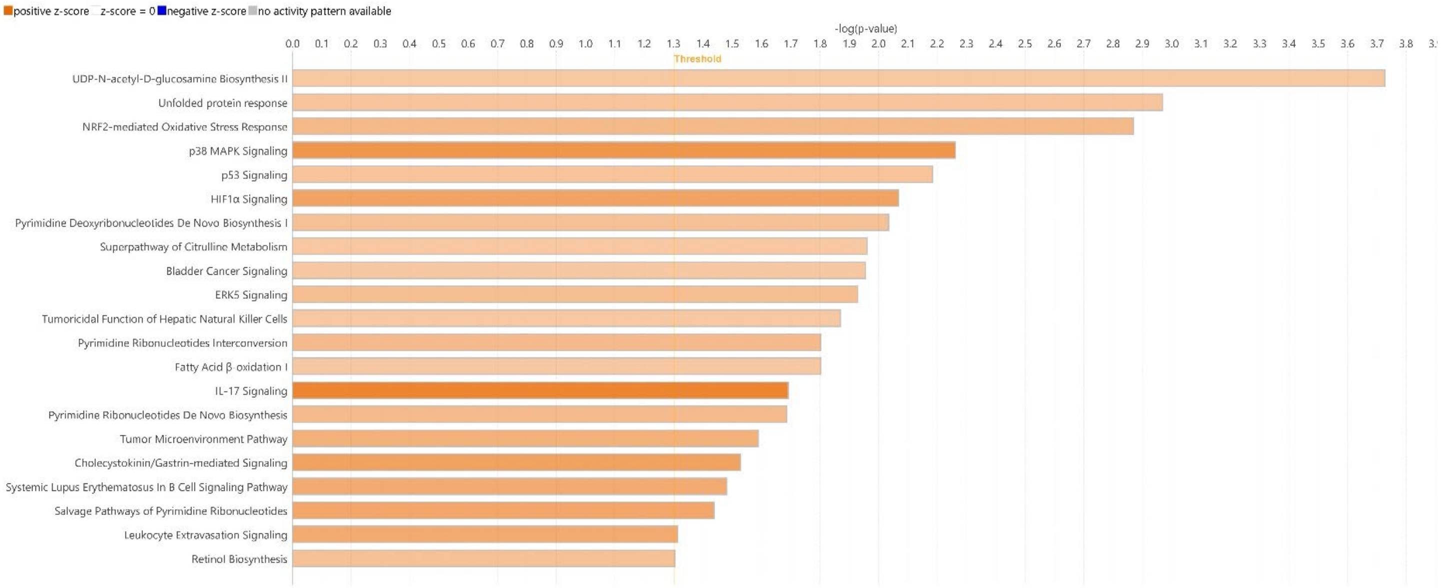


© 2000-2023 QIAGEN. All rights reserved.

**Supplemental Figure 9. Indole upregulates transcriptional pathways in human colon B cells.** LPMCs from healthy human colon tissue were stimulated with 1 mM indole or vehicle for 4hr followed by RNA was isolated from flow-sorted CD19+ B cells for RNAseq. Differentially expressed pathways (indole vs vehicle) were identified with Ingenuity Pathway Analysis for CD19+ B cells. N=5 paired samples.

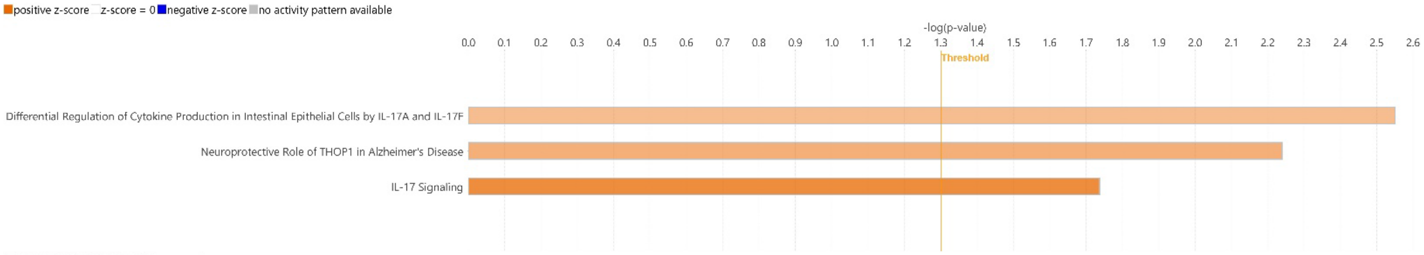


© 2000-2023 QIAGEN. All rights reserved.

**Supplemental Figure 10. Indole upregulates transcriptional pathways in human colon T cells.** LPMCs from healthy human colon tissue were stimulated with 1 mM indole or vehicle for 4hr followed by RNA was isolated from flow-sorted CD3+ T cells for RNAseq. Differentially expressed pathways (indole vs vehicle) were identified with Ingenuity Pathway Analysis for CD3+ T cells. N=5 paired samples.
